# Two broadly conserved families of polyprenyl-phosphate transporters

**DOI:** 10.1101/2022.02.03.479048

**Authors:** Ian J. Roney, David Z. Rudner

## Abstract

Peptidoglycan and virtually all surface glycopolymers in bacteria are built in the cytoplasm on the lipid carrier undecaprenyl-phosphate (UndP). These UndP-linked precursors are transported across the membrane and polymerized or directly transferred to surface polymers, lipids, or proteins. UndP is then flipped to regenerate the pool of cytoplasmic-facing UndP. The identity of the flippase that catalyzes transport has eluded identification for decades. Here, using the antibiotic amphomycin that targets UndP, we discovered two broadly conserved families that catalyze UndP recycling. One (UptA) is a member of the DedA superfamily; the other (PopT) contains the domain DUF368. We show that family members from gram-positive and gram-negative bacteria catalyze UndP transport in *Bacillus subtilis*. Inhibitors of these flippases could potentiate the current arsenal of cell envelope-targeting antibiotics.

**One Sentence Summary:** We define two transporter families that recycle the universal lipid carrier in surface glycopolymer biogenesis.

## Main Text

Polyprenyl-phosphates are universal carriers of sugars and glycopolymers across membranes in all domains of life (*1–4*). In bacteria, the 55-carbon isoprenoid undecaprenyl-phosphate (UndP) is used to transport most glycopolymers across the cytoplasmic membrane including peptidoglycan (PG) precursors, O-antigens of lipopolysaccharide (LPS), teichoic acids, and capsule. UndP also ferries sugars and oligosaccharides across the membrane that are used to glycosylate lipid A of LPS, teichoic acids, and surface proteins (**Fig. S1**). In eukaryotes and archaea the polyprenyl-phosphate, dolichol-phosphate (DolP) transports oligosaccharides across the membranes of the endoplasmic reticulum and the archaeal cytoplasmic membrane, respectively (*1, 2*). These DolP-linked sugars are then used to decorate membrane-anchored and secreted proteins. In all cases, the lipid carrier must be recycled via transport across the cytoplasmic membrane. The flippases that catalyze polyprenyl-phosphate transport are among the last unknown enzymes in these pathways.

Bacterial cell wall biogenesis, the target of some of the most effective antibiotics, exemplifies these glycopolymer synthesis pathways (**Fig. 1**) (*5*). A series of cytoplasmic enzymatic steps generate the PG precursor lipid II, a disaccharide pentapeptide linked to undecaprenyl-pyrophosphate (UndPP). Lipid II is then flipped to the outer leaflet of the cytoplasmic membrane where the muropeptide is polymerized and crosslinked into the cell wall meshwork. The UndPP product is dephosphorylated and UndP is flipped to the inner leaflet to complete the lipid II cycle (**Fig. 1**). During exponential growth, a bacterium contains ∼10^5^ UndP molecules that are thought to ferry a similar number of PG precursors across the cytoplasmic membrane every minute, suggesting that each lipid carrier is reused multiple times per cell cycle for PG synthesis alone (*6, 7*). *De novo* synthesis of UndP maintains the pool size during growth while recycling is the primary source for surface polymer biogenesis. Here, we define two broadly conserved families of flippases that catalyze UndP transport in bacteria and could function to recycle DolP in eukaryotes and archaea.

**Figure 1.**
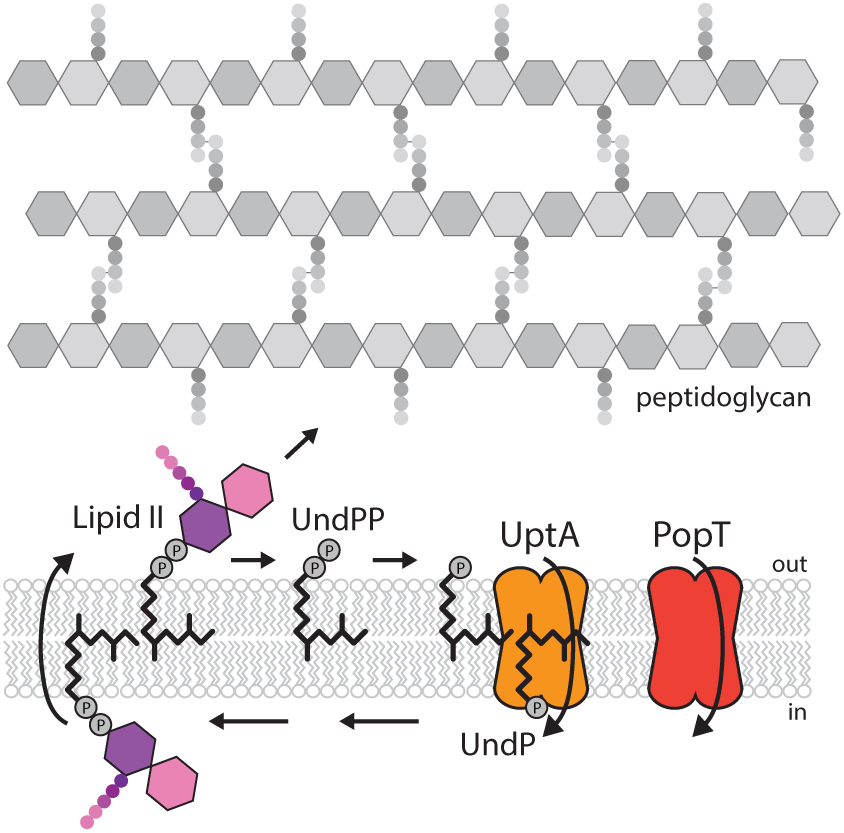
UptA and PopT complete the lipid II cycle. Schematic of the lipid II cycle in peptidoglycan synthesis highlighting the role of UptA and PopT in catalyzing the transport of undecaprenyl-phosphate.

To screen for potential UndP flippases, we took advantage of the antibiotic amphomycin and its derivative MX2401 (*8–10*). These cyclic lipopeptides bind the surface-exposed phosphate on UndP preventing its recycling and thus inhibiting cell wall synthesis. We reasoned that over-expression of an UndP transporter would provide resistance to these antibiotics. We generated a *Bacillus subtilis* transposon library with a strong outward facing promoter (*11*) and plated the library on LB agar supplemented with MX2401 at concentrations above the minimum inhibitory concentration (MIC). The colonies were pooled and the transposon insertion sites mapped by transposon-sequencing (Tn-seq). Tn insertions were significantly over-represented at a single region of the genome and in all cases the outward-facing promoter faced the uncharacterized gene *yngC* (**Fig. 2A**). Over-expression of *yngC in B. subtilis* increased the MIC of MX2401 by >16-fold (**Fig. 2B**), validating our findings. Furthermore, deletion of *yngC* reduced the MIC >4-fold (**Fig. 2B**). These data argue that YngC functions in UndP transport.

**Figure 2.**
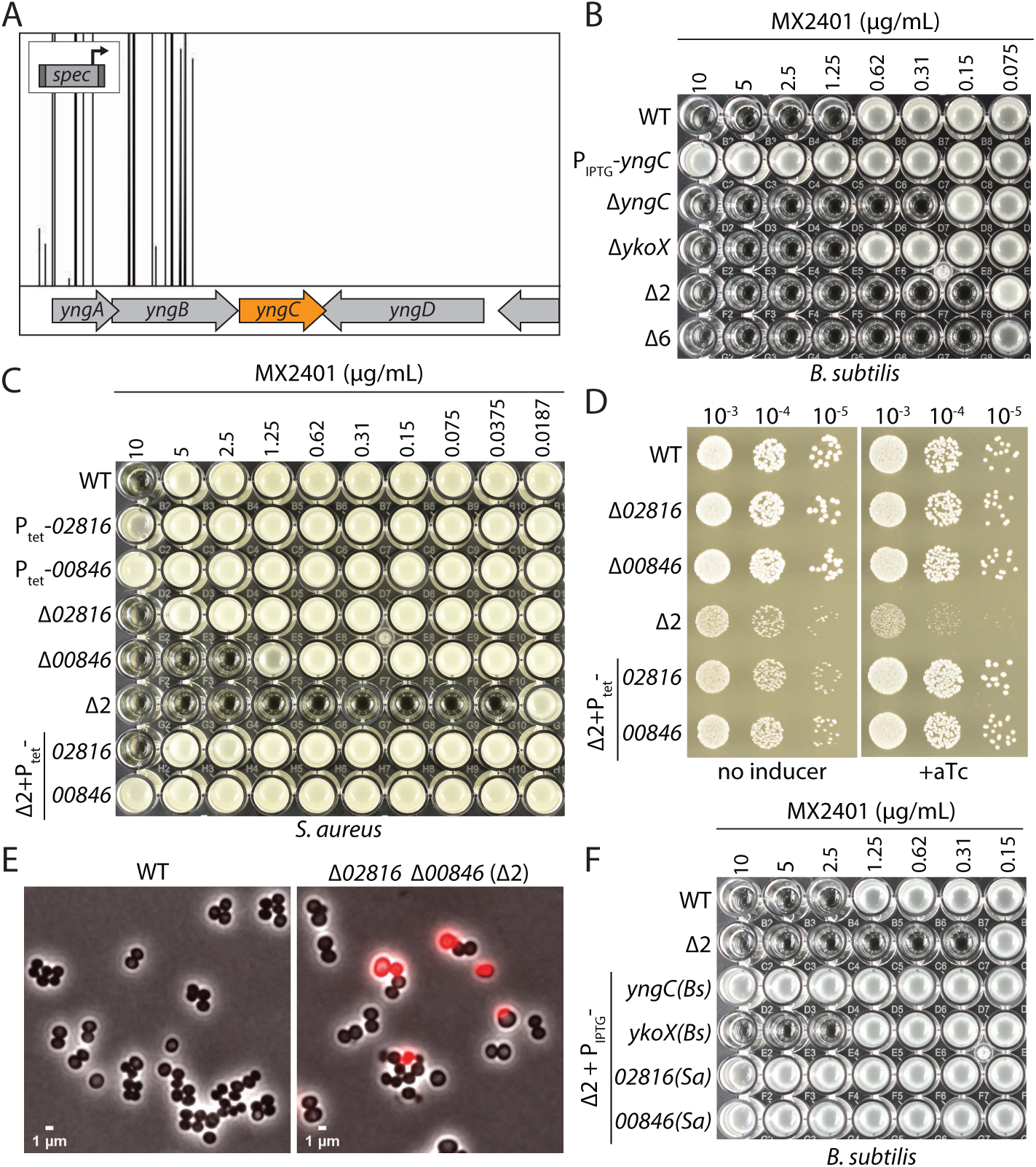
DedA and DUF368 family members provide resistance to UndP targeting antibiotics. **(A)** A transposon-sequencing screen for MX2401 resistance mutants identifies *yngC*. Transposon insertion profile at the indicated *B. subtilis* genomic region is shown. Each vertical line indicates an insertion site; its height reflects the number of sequencing reads at this position (maximum height ≥5,000). All transposons in this region were inserted with the outward-facing promoter facing *yngC* as indicated in the schematic. Most insertion sites had >20,000 reads but the window size was scaled to 5,000 to highlight the absence of reads in the neighboring genes. **(B)** Minimum inhibitory concentration (MIC) assay of the indicated *B. subtilis* strains. **(C)** MIC assay of the indicated *S. aureus* strains. **(D)** Spot-dilution assays of the indicated *S. aureus* strains on TSA with or without 100 ng/mL anhydrotetracycline (+aTc). **(E)** Representative images of wild-type and *Δ00846 Δ02816 S. aureus* cells. An overly of phase-contrast and fluorescent images of propidium iodide (PI)-stained cells are shown. **(F)** MIC assays of the indicated *B. subtilis* strains over-expressing DedA and DUF368 family members from *B. subtilis* and *S. aureus*.

YngC is a member of the DedA superfamily, a broadly conserved but poorly understood family of membrane proteins found in all domains of life (*12*). The AlphaFold2-predicted structure of YngC resembles membrane transporters with membrane re-entrant loops on either side of the lipid bilayer (*13*) (**Fig. S2A**). A conserved pair of arginines in one of these loops is required for MX2401 resistance (**Fig S2B**) and could function in phosphate binding. Interestingly, one of *yngC*’s promoters is regulated by the extracytoplasmic function (ECF) sigma factor σ^M^ (*14*) (**Fig. S3**). This transcription factor is induced under conditions that trap UndP-linked intermediates leading to a reduction in the cellular pools of the lipid carrier (*15*). Consistent with a role for σ^M^ in regenerating this pool, a *sigM* mutant or a deletion of the σ^M^-binding site in the *yngC* promoter sensitized *B. subtilis* to MX2401 (**Fig S3**). Finally, we found that reducing *de novo* synthesis of UndP also sensitized *B. subtilis* to loss of *yngC* (**Fig. S4**), providing further support for the idea that YngC recycles the lipid carrier.

*B. subtilis* encodes five additional DedA paralogs. Individual deletions in these genes did not affect the MIC of MX2401 (**Fig. S5**). A deletion of one of the paralogs, *ykoX*, modestly reduced the MIC of the Δ*yngC* mutant and a strain lacking all six paralogs phenocopied the Δ*yngC ΔykoX* double mutant (**Fig 2B**). Thus, YngC is likely to be the principal *B. subtilis* DedA family member involved in UndP recycling. We note that over-expression of YkoX in the Δ*yngC* mutant provided some resistance to MX2401 (**Fig. 2F and S6**). These data raise the possibility that YkoX transports a distinct anionic lipid but can act on UndP when over-expressed.

Tn-seq screens in *Staphylococcus aureus* to profile the phenotypic responsiveness to 32 antibiotics identified transposon insertions upstream of two genes (*SAOUHSC_02816* and *00846*) when the Tn library was challenged with sub-inhibitory doses of amphomycin (*16*) (**Fig. S7**). *02816* encodes a DedA family member and *00846* encodes a protein with a domain of unknown function DUF368. We validated these uncharacterized hits and found that over-expression of either protein in *S. aureus* increased the MIC of MX2401 (**Fig. 2C**). Deletion of *02816* did not impact the MIC of MX2401 while deletion of *00846* reduced it by 4-fold (**Fig. 2C**). Strikingly, the MIC of the double mutant was ∼256-fold lower than wild-type (**Fig. 2C**). Furthermore, the double mutant was growth-impaired, displayed cell size variability, and 10% of the cells had membrane permeability defects, consistent with impaired envelope assembly (**Fig 2E and S8**). Finally, over-expression of either *S. aureus* gene in *B. subtilis* lacking *yngC* and *ykoX* provided resistance to MX2401 (**Fig 2F**). Since *B. subtilis* lacks a DUF368 homolog, these data argue that the *S. aureus* DUF368 protein functions as an UndP transporter rather than a regulator or co-factor. DUF368 is a polytopic membrane protein that is widely conserved in bacteria and archaea (**Fig. S9**). The AlphaFold2-predicted structure of *S. aureus* DUF368 resembles canonical membrane transporters with 2-fold inverted symmetry and membrane re-entrant loops on either side of the lipid bilayer (*13, 17*) (**Fig. S10**). The predicted gap between transmembrane helices 4 and 11 could allow the polyprenyl tail to remain in the lipid bilayer during transport of the phosphate head group, akin to the membrane-spanning groove in the lipid scramblases in the eukaryotic TMEM16 family (*18, 19*). Based on the findings presented thus far and those below, we have renamed the DedA superfamily members encoded by *yngC* and *02816* UptA (<u>U</u>nd<u>P t</u>ransporter <u>A</u>) and DUF368 proteins PopT (<u>po</u>lyprenyl-<u>p</u>hosphate transporter).

Gene neighborhood analysis (*20*) provides additional support for a role of these two protein families in UndP transport. In a subset of bacterial genomes, genes encoding a DedA superfamily member and a lipid phosphatase of the PAP2 family (*21*) are adjacent to each other; in others the two genes are fused (**Fig. 3A and S11**). Some PAP2 homologs, like *B. subtilis* BcrC and *E. coli* PgpB, are undecaprenyl-pyrophosphate phosphatases that convert surface-exposed UndPP to UndP prior to their transport to the cytoplasmic face of the membrane (*22*). Similarly, in other bacterial genomes a *dedA* gene is present adjacent to *uppP* that encodes a second family of UndPP phosphatases (*23*) (**Fig S12**). Accordingly, if UptA transports UndP then these adjacent or fused genes would encode enzymes that catalyze sequential steps in the lipid II cycle (**Fig 1A**). In the case of PopT, most archaeal genomes contain a cluster of genes required for surface protein glycosylation (*24*). A DUF368 containing gene is also present in many archaeal genomes and 37% of the time is found in these clusters, often adjacent to an oligosaccharyltransferase of the *aglB* family (**Fig 3B and S13**). The transfer of the lipid-linked oligosaccharide onto a surface protein by AglB liberates DolP that could then be recycled by the PopT homolog. Finally, gram-positive bacteria encode enzymes involved in glycosylating surface polymers and a DedA superfamily member is sometimes encoded in an operon with these enzymes. *B. subtilis* provides a prototypical example (**Fig. S14A**). In this bacterium, a transcription factor controls the expression two operons that are thought to be involved in glycosylation of lipoteichoic acids (*25*). Four of the genes in these operons encode the full set of enzymes required to glycosylate this surface polymer. The fifth encodes UptA, which would complete the cycle in this sugar modification pathway (**Fig. S14B**). Thus, gene neighborhood analysis suggests UptA and PopT family members are broadly conserved polyprenyl-phosphate transporters.

**Figure 3.**
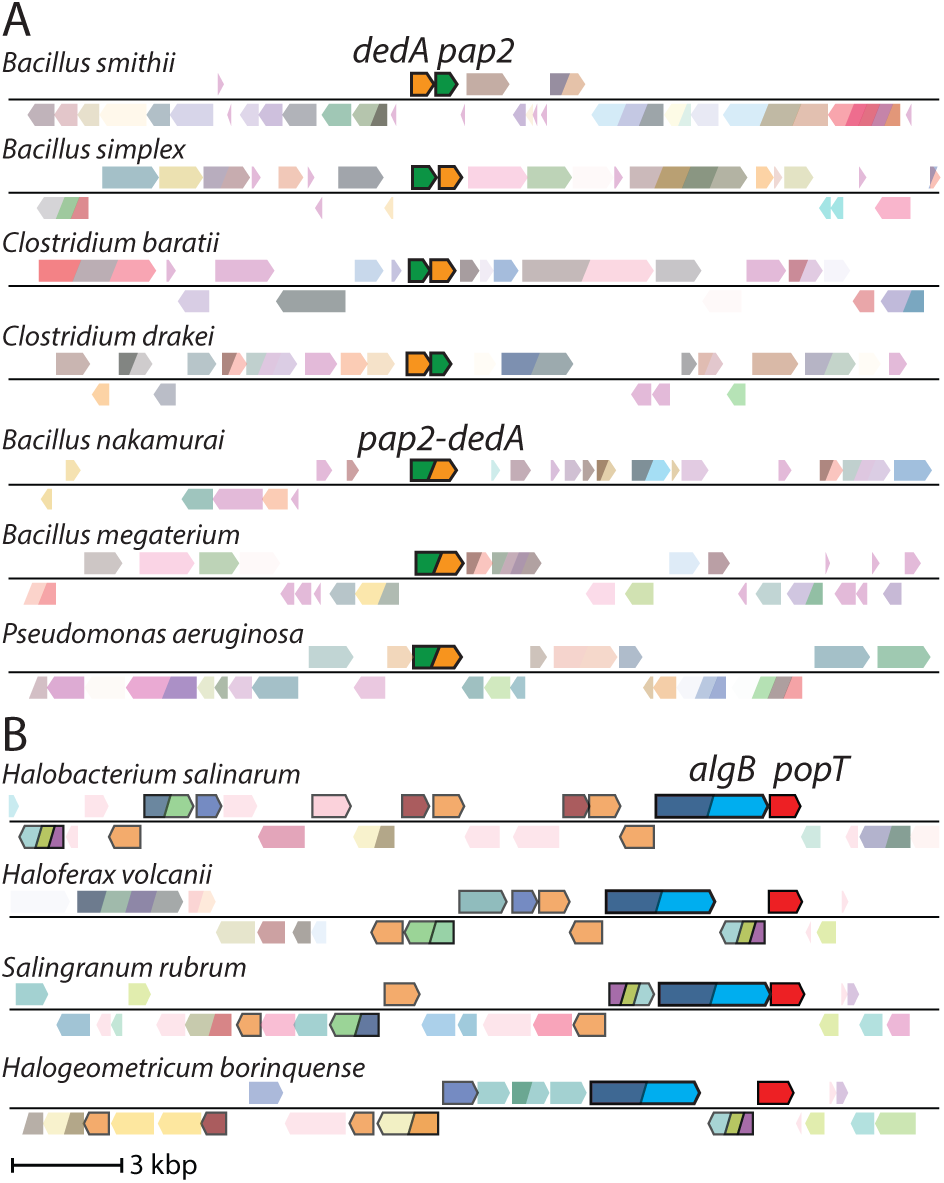
Gene neighborhood analyses for DedA and DUF368 family members. Representative genomic neighborhood diagrams highlight the synteny or fusion of a *dedA* gene with a gene encoding a PAP2 lipid phosphatase **(A)** and the synteny of *duf368* with genes involved in archaeal N-linked protein glycosylation pathways **(B)**. ∼1.3% of all *dedA* genes are fused to *pap2* genes. The gene encoding the oligosaccharyltransferase AglB is shown in blue and other genes in the glycosylation pathway are highlighted by black borders. 37% of *duf368* genes are present in gene clusters encoding protein glycosylation pathways.

To directly test whether UptA and PopT family members function in UndP recycling, we fluorescently labeled MX2401 and used it to visualize surface-exposed UndP. Wild-type *B. subtilis* cells expressing cytoplasmic blue fluorescent protein (BFP) and the Δ*uptA* Δ*ykoX* double mutant expressing mCherry were mixed and then labeled with MX2401-FL and visualized by fluorescence microscopy. As can been seen in Figure 4A, wild-type cells had faint MX2401-FL signal while cells lacking the two DedA paralogs were more strongly stained along the cytoplasmic membrane. Importantly, the MX2401-FL signal in the *B. subtilis* double mutant was dramatically reduced when UptA^Bs^, UptA^Sa^, or PopT^Sa^ were over-expressed (**Fig. 4A and S15**). In an analogous experiment, we labeled wild-type *S. aureus* and the Δ*uptA* Δ*popT* double mutant with spectrally distinct Fluorescent D-amino acids (FDAAs) (*26*) and then mixed the two cultures and stained with MX2401-FL. The double mutant had strong MX2401-FL fluorescence while the signal was barely detectable in wild-type (**Fig. 4A and S15**).

**Figure 4.**
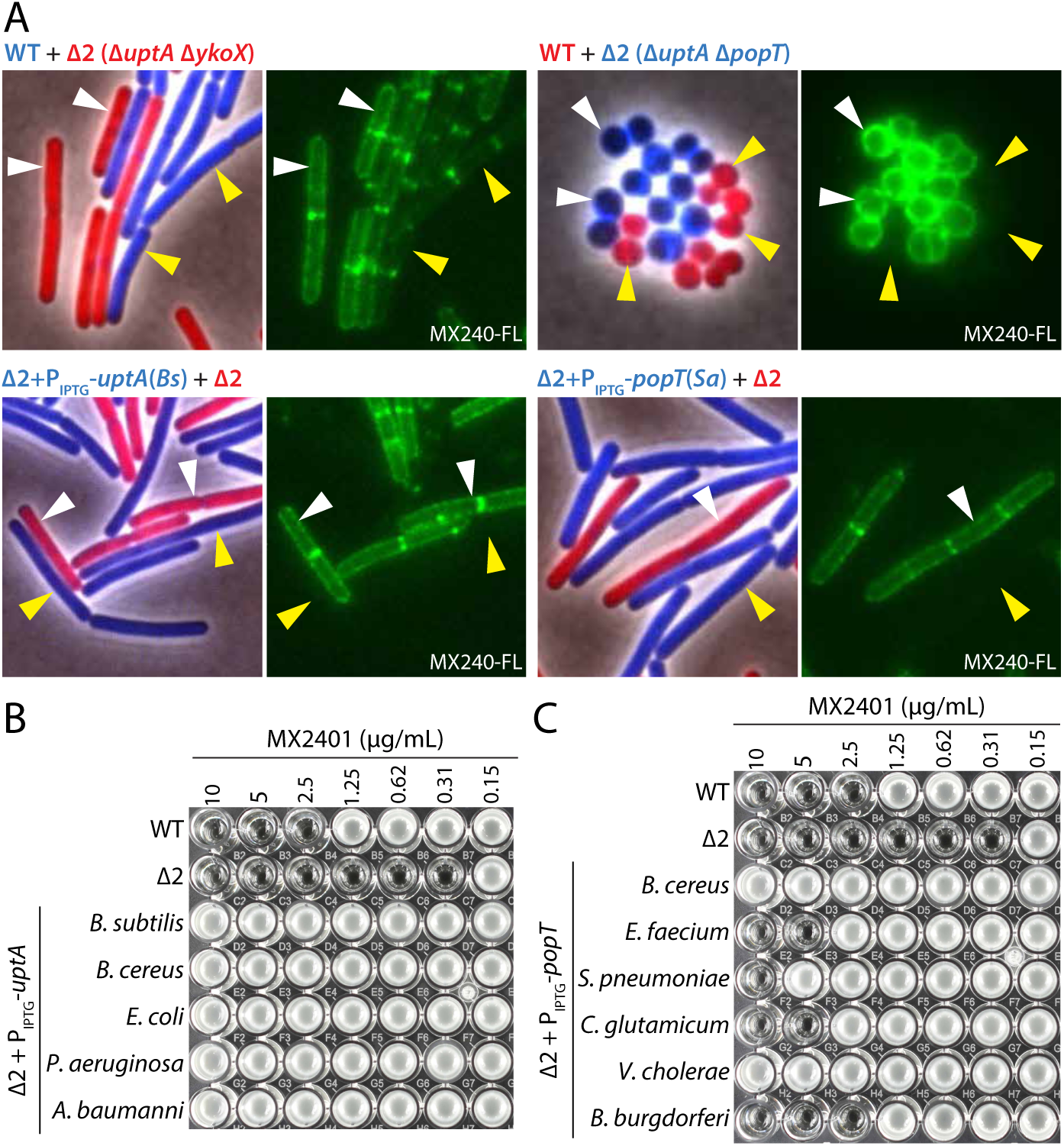
UptA and PopT are broadly conserved UndP transporters. **(A)** Representative microscopy images of the indicated *B. subtilis* or *S. aureus* strains. Two strains expressing different fluorescent proteins (*B. subtilis*) or labeled with different fluorescent D-amino acids (*S. aureus*) were mixed and then stained with fluorescent MX2401 (MX2401-FL). The left panels show overlays of phase-contrast and fluorescent images in the red and blue channels to distinguish the two strains. The right panels show MX2401-FL staining. Yellow carets highlight wild-type cells or cells over-expressing UptA(*Bs*) or PopT(*Sa*). White carets highlight cells lacking the UndP transporters. **(B)** MIC assays of *B. subtilis* strains over-expressing UptA homologs from the indicated bacteria. **(C)** MIC assays of *B. subtilis* strains over-expressing PopT homologs from the indicated bacteria.

*E. coli* encodes eight DedA superfamily members (*12*). The mutants have pleiotropic phenotypes that include defects in cell division, morphology, alkaline tolerance, membrane potential, and antibiotic resistance. The proteins have been hypothesized to function as proton-dependent transporters that help maintain the proton motive force (*27*). To investigate whether any of these factors catalyze UndP transport, we expressed each in *B. subtilis* and tested for MX2401 resistance. Although several paralogs increased the MIC of MX2401 of the *B. subtilis* Δ*uptA* Δ*ykoX* mutant by ∼2-4-fold, only one conferred high-level resistance (**Fig. 4B and S16A**). This DedA paralog is the namesake of the superfamily and is most similar to UptA^Bs^ and UptA^Sa^ (**Fig. S2A and Table S1A**). We extended this cross complementation analysis to a diverse collection of UptA and PopT homologs. The DedA superfamily members tested were chosen based on their similarity to the validated UptA family members. In addition to UptA^Sa^ and UptA^Ec^, homologs from *Acinetobacter baumannii*, *Pseudomonas aeruginosa*, and *Bacillus cereus* conferred MX2401 resistance to the *B. subtilis* double mutant (**Fig. 4B**). Similarly, expression of PopT homologs from *Bacillus cereus*, *Streptococcus pneumoniae*, *Vibrio cholerae*, *Borrelia burgdorferi*, and *Corynebacterium glutamicum* increased the MIC of MX2401 (**Fig 4C**). Importantly, expression of these proteins in *B. subtilis* also reduced surface-exposed UndP as assayed by MX2401-FL (**Fig. S17 and S18**).

We conclude that UptA and PopT family members catalyze the transport of UndP, the final step in virtually all cell envelope biogenesis and surface modification pathways in both gram-positive and gram-negative bacteria. Their discovery completes the parts list for many of these intensively studied pathways and represents a new set of targets to potentiate the current arsenal of antibiotics.

The mechanisms of UndP transport by UptA and PopT are currently unknown but recent in vitro reconstitution experiments of two eukaryotic DedA superfamily members TMEM41B and VMP1 provide possible insight (*28*). Both proteins were found to redistribute fluorescently labeled phospholipids with diverse headgroups from the luminal to the exposed face of proteoliposomes. Lipid transport was energy independent suggesting these transporters function as lipid scramblases. These data lead us to propose that both UptA and PopT family members function as UndP scramblases rather than flippases. In the context of this model transfer of sugars onto the lipid carrier on the cytoplasmic face of the membrane would result in unidirectional flux across the lipid bilayer. Interestingly, depletion of either TMEM41B or VMP1 in tissue culture cells led to defects in the sorting of cholesterol, phosphatidylserine, and phosphatidylcholine providing support for a role in lipid transport in vivo (*28, 29*). The analysis of UptA family members presented here strengthens these conclusions and suggests that other bacterial DedA superfamily members function in lipid transport. Based on our analysis of the *B. subtilis* DedA paralogs, we hypothesize that in vivo these transporters have more narrow substrate specificity and act on distinct lipids. Furthermore, it is tempting to speculate that a DedA superfamily member recycles DolP in eukaryotes while PopT homologs transport DolP in archaea.

Other than an increase in surface-exposed UndP, the *B. subtilis* Δ*uptA* Δ*ykoX* double mutant had no discernable growth or morphological defects. The *S. aureus* Δ*uptA* Δ*popT* mutant was growth-impaired and had membrane permeability defects but was viable. These data raise the possibility that a third family of UndP transporters exist or that UndP flipping can occur spontaneously albeit inefficiently. We favor the latter model because a screen using a transposon with an even stronger outward-facing promoter did not identify additional hits when the *B. subtilis* Δ*uptA* Δ*ykoX* double mutant was challenged with MX2401. Similarly, Tn insertions were only enriched upstream of *uptA* and *popT* in the analogous screen in *S. aureus* (*16*). Using over-expression libraries from *B. subtilis*, *S. aureus*, and *B. cereus*, we only identified UptA and PopT homologs that could confer MX2401 resistance in the *B. subtilis* Δ*uptA* Δ*ykoX* mutant. Finally, no genes were identified as synthetic lethal with the single or double *B. subtilis* mutants. UppP has been hypothesized to function as both an UndPP phosphatase and an UndP transporter (*3, 30*) but over-expression or deletion of *uppP* or *bcrC* in *B. subtilis* had not impact on the MIC of MX2401 (**Fig. S16B**) and neither gene was identified as a hit in our Tn-seq screen nor the one performed in *S. aureus* (**Fig. S6 and S7**). We therefore propose that UndP can spontaneously flip/flop in the cytoplasmic membrane and hypothesize that the extent of protonation of the phosphate headgroup influences efficiency and could explain the defect in alkaline tolerance in the *E. coli* mutants (*31*) and the difference in mutant phenotypes in *B. subtilis* and *S. aureus*.

Although not essential, the DedA superfamily member that is most similar to *E. coli* DedA (UptA^Ec^) in *Neisseria meningitidis, Klebsiella pneumoniae* and *Burkholderia thailandensis* (**Table S1B**) is required for intrinsic resistance to the last-resort antibiotic colistin (*27, 32, 33*). Loss-of-function mutations in the associated genes in these gram-negative pathogens confer colistin-sensitivity and in some cases it has been shown that expression of UptA^Ec^ can reverse this sensitivity, implicating UndP transport. Interestingly, colistin resistance is mediated by aminoarabinose (Ara4N) modification to lipid A of LPS, which reduces its affinity for polycationic antibiotics(*34*). The Ara4N modification is built on UndP in the cytoplasm, flipped to the periplasm and then transferred to lipid A. Mutations in these DedA superfamily members result in low levels of Ara4N-modified lipid A (*27*) and we hypothesize that the reduction in UndP recycling in these mutants accounts for the decreased modification. UptA homologs in *Klebsiella pneumoniae* and *A. baumannii* are also required for resistance to complement-mediated killing and *K. pneumoniae* UptA is additionally required for neutrophil evasion in a mouse model for lung colonization (*35–37*). Long O-antigen chains and a thick capsule layer, both glycopolymers that are built on UndP, are primary defense mechanisms of these pathogens. Thus small molecules that target UptA and PopT transporters could potentiate both last resort antibiotics and host defense mediated killing.

## Acknowledgements

We thank all members of the Bernhardt-Rudner super-group for helpful advice, discussions, and encouragement, Tom Bernhardt, Suzanne Walker, Andrew Kruse for insights and advice, Susan Farmer and Bob Hancock for MX2401, the MicRoN core for advice on microscopy, and Matt Waldor for coordinating submission. Support for this work comes from the National Institute of Health Grants GM086466, GM127399, U19 AI109764 (D.Z.R.).

## Supplemental Figure Legends

**Figure S1.**
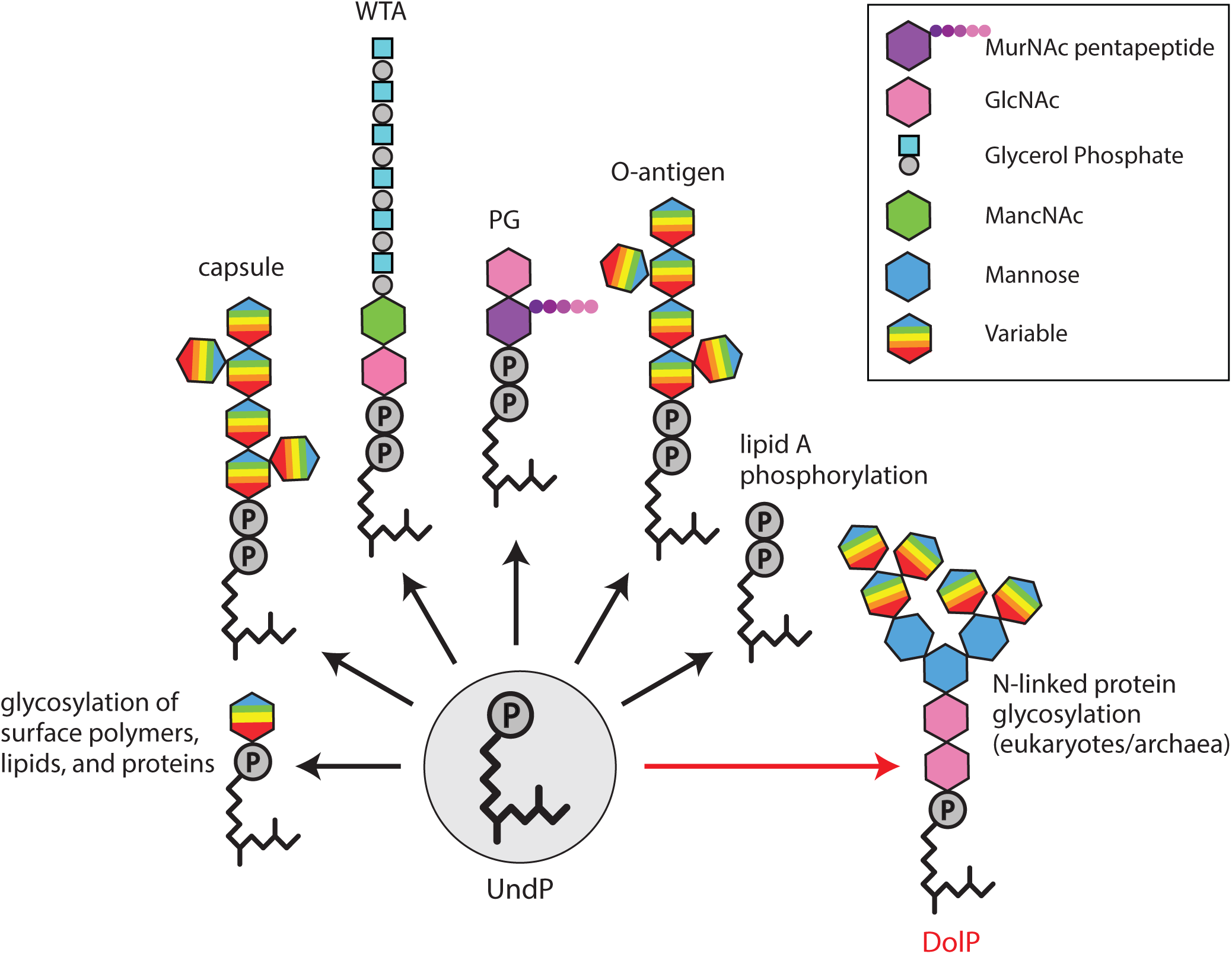
Polyprenyl-phosphate lipids are universal transporters of surface glycopolymers. Schematic illustrating the glycopolymers and sugar modifications that are assembled on undecaprenyl-phosphate (UndP) in bacteria and on dolichol-phosphate (DolP) in archaea and eukaryotes. Although an oligosaccharide carried on DolP is shown, some eukaryotes use the pyrophosphate carrier (DolPP).

**Figure S2.**
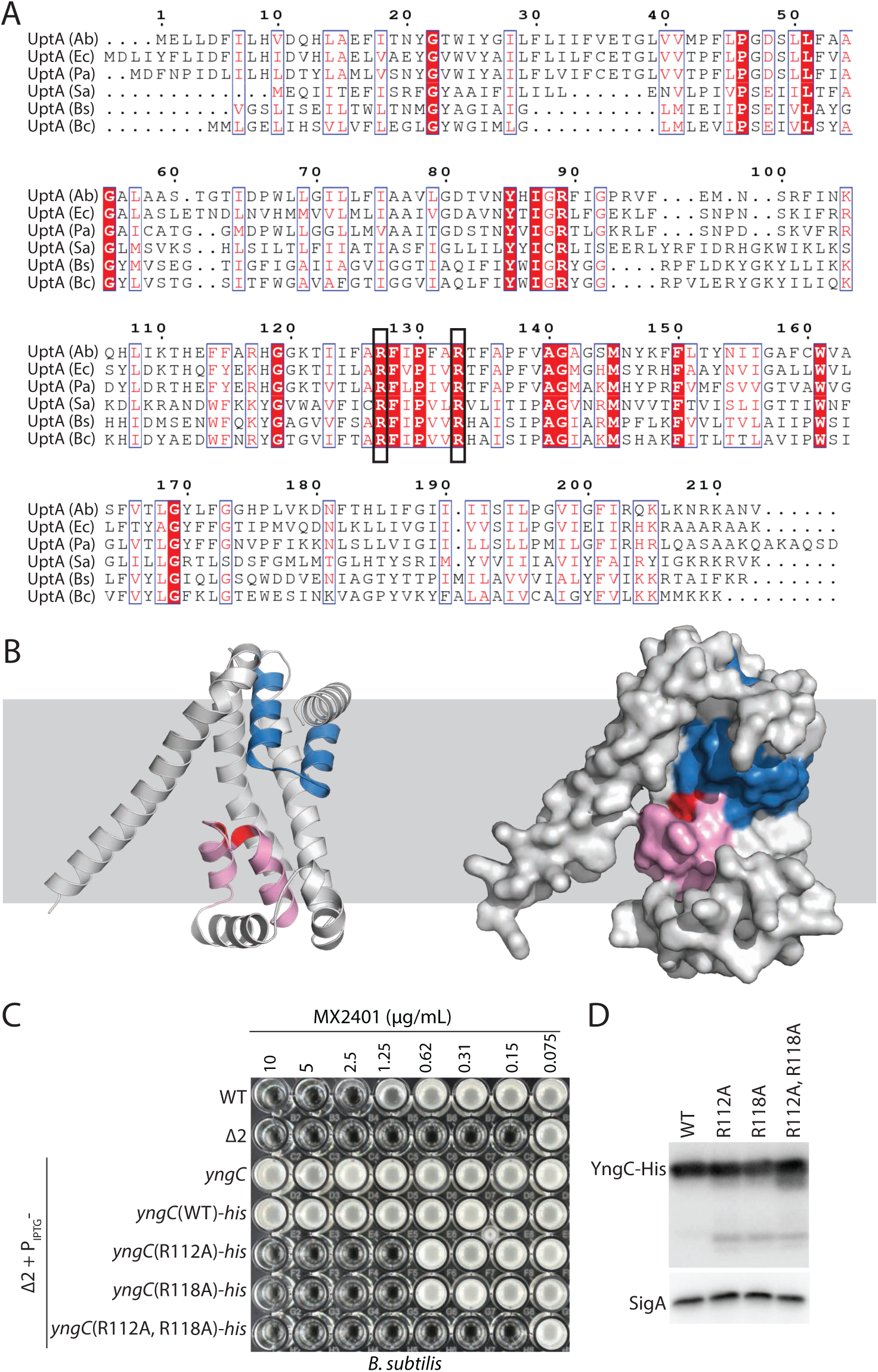
The predicted structure of YngC has features of membrane transporters. **(A)** Multiple sequence alignment of DedA (UptA) homologs that provide MX2401 resistance. Two highly conserved arginines are boxed. **(B)** Structural models of YngC as predicted by AlphaFold2. Membrane re-entrant helices that are commonly found in membrane embedded transporters (*13*) are highlighted in red and blue. **(C)** Minimum inhibitory concentration (MIC) assays of the indicated *B. subtilis* strains with point mutations in the conserved arginines in *yngC*. **(D)** Immunoblot analysis of YngC-His levels using anti-His antibodies of the strains in (C). SigA serves as a control for loading.

**Figure S3.**
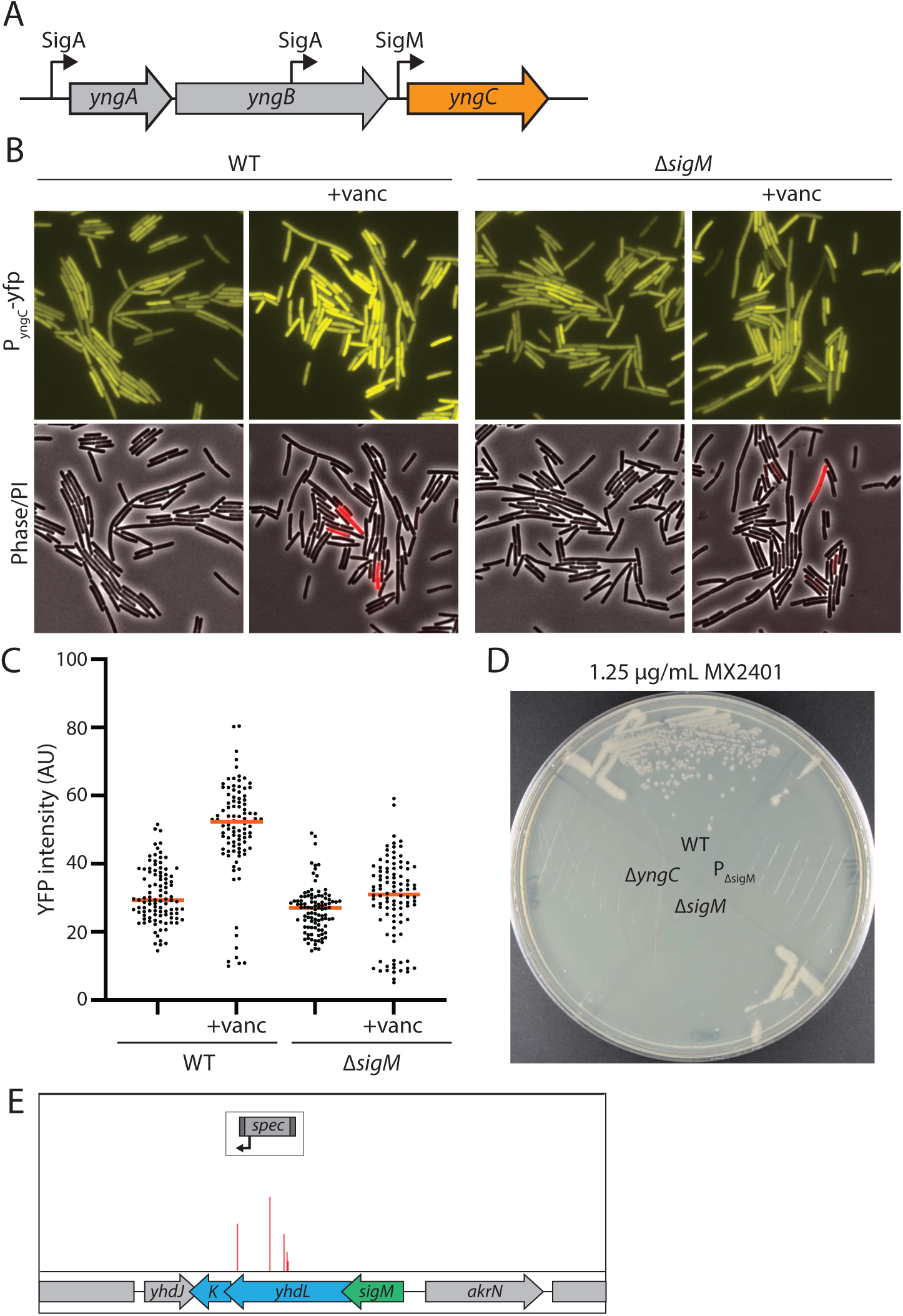
σ^M^ transcription of *yngC* contributes to MX2401 resistance. **(A)** Schematic of the *yngABC* operon highlighting the three promoters that regulate *yngC* expression. Two promoters are recognized by sigma factor A (σ^A^) and the third is recognized by the ECF sigma factor M (σ^M^). **(B)** Representative fluorescent images of cells harboring a transcriptional fusion of the *yngC* σ^M^ promoter to *yfp*. YFP fluorescence increases in cells exposed to vancomycin for 30 minutes, a condition that activates σ^M^. The transcriptional reporter is not induced by vancomycin in a *ΔsigM* mutant. **(C)** Quantification of YFP fluorescence from the strains and conditions in (B). **(D)** Streaks of the indicated strains on LB agar supplemented with 1.25 µg/mL MX2401. P_Δ*sigM*_ contains a deletion of the σ^M^ promoter of *yngC*. **(E)** In the Tn-seq screen for MX2401 resistance mutants, a small number of insertions mapped to the genes (*yhdL* and *yhdK*) encoding the anti-σ^M^ factors, consistent with increased σ^M^-dependent transcription of *yngC* providing MX2401 resistance. Transposon insertion profile at the indicated *B. subtilis* genomic region is shown. Each vertical line indicates an insertion site; its height reflects the number of sequencing reads at this position (maximum height ≥5,000). The transposon insertion site with the maximum number of reads in this region had 2,600 reads. For comparison, most of the insertion sites adjacent to *yngC* had >20,000 reads.

**Figure S4.**
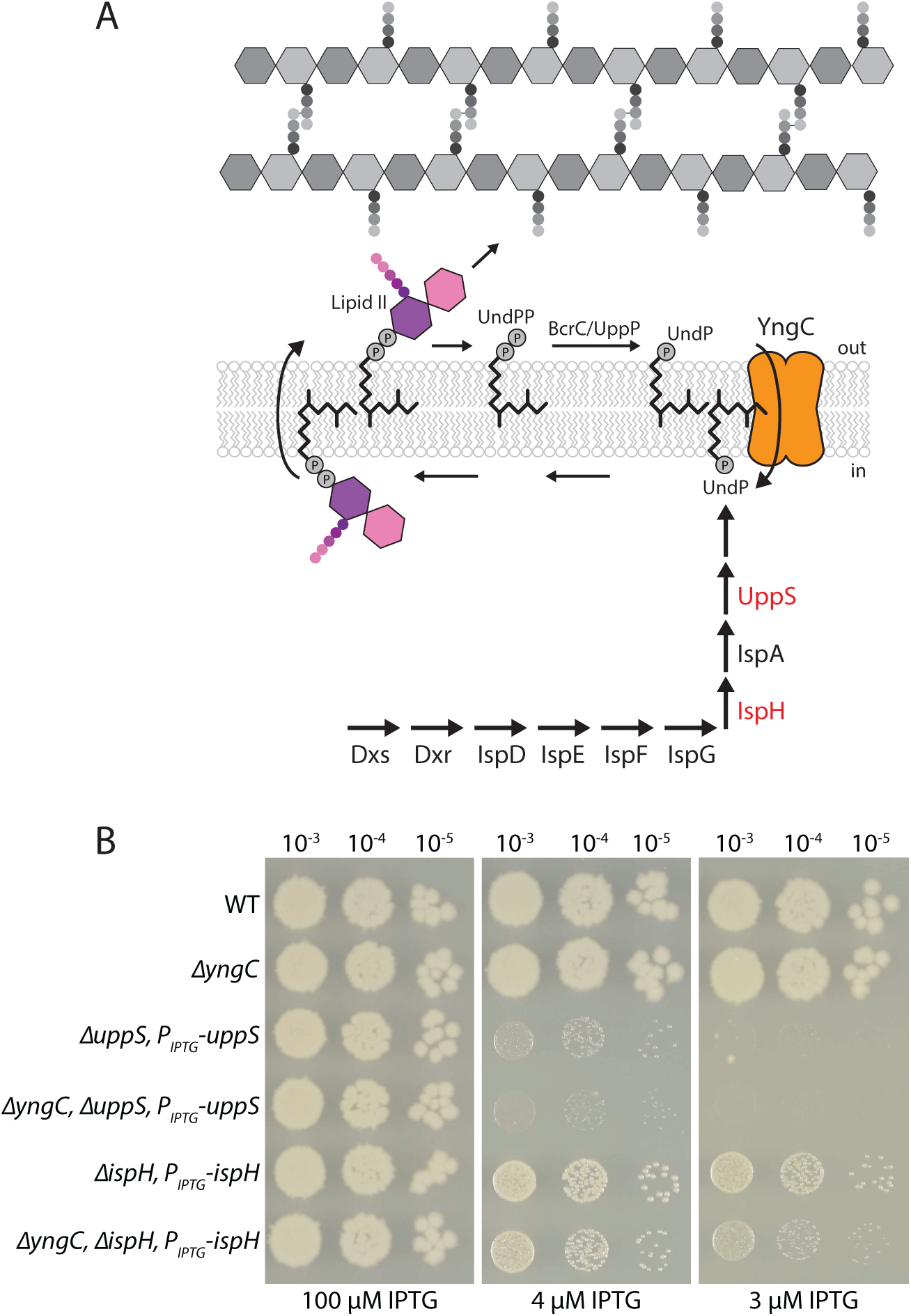
Δ*yngC* sensitizes *B. subtilis* to reduced levels of UndP synthesis. **(A)** Schematic of the UndP synthesis pathway illustrating the two sources of UndP for the lipid II pathway, *de novo* synthesis and recycling. **(B)** Spot-dilution assays of the indicated *B. subtilis* strains with IPTG-regulated alleles of *uppS* or *ispH* on LB agar supplemented with the indicated concentrations of IPTG.

**Figure S5.**
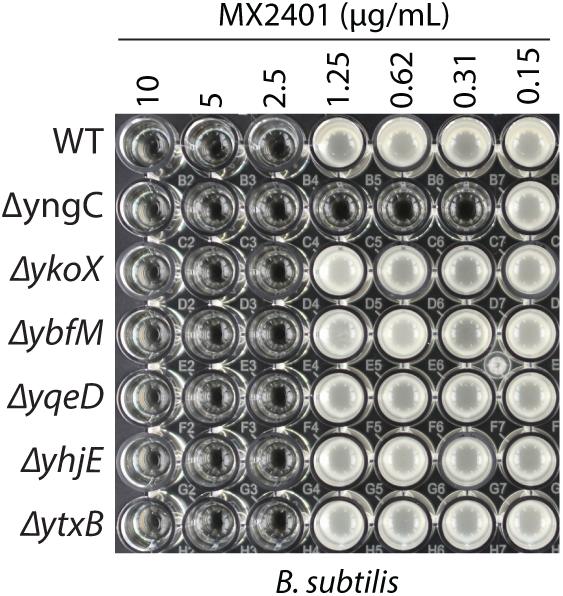
YngC is the principal DedA family member that provides resistance to MX2401. Minimum inhibitory concentration (MIC) of MX2401 in the indicated *B. subtilis* strains, each lacks one of the six DedA paralogs.

**Figure S6.**
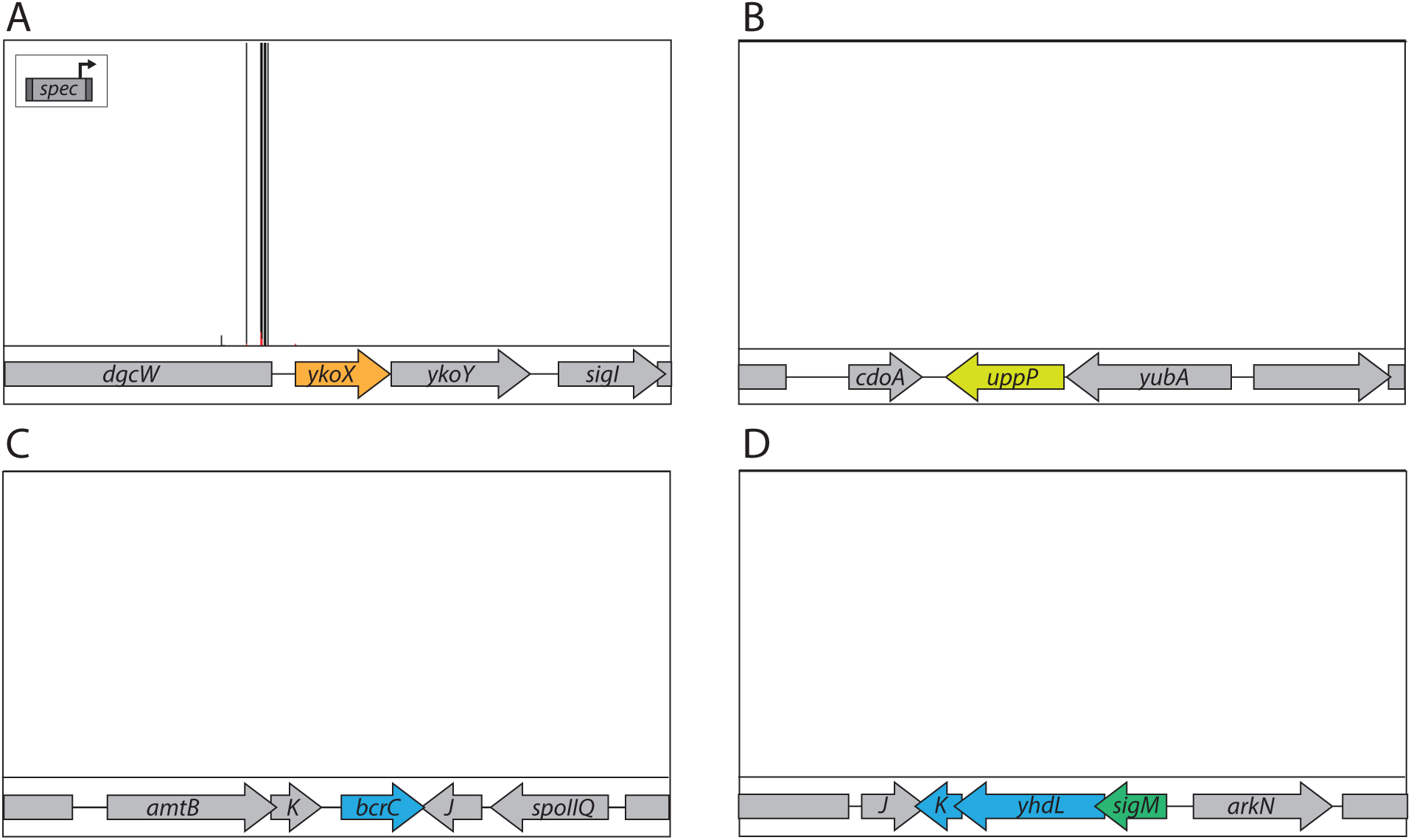
Transposon insertions that increase transcription of *ykoX* provide resistance to MX2401 in a *B. subtilis* mutant lacking YngC. A *B. subtilis* Δ*yngC* mutant strain was transposon-mutagenized with a transposon carrying a strong outward facing P*_pen_* promoter (insert). The library was plated on LB agar supplemented with 0.3 µg/mL MX2401 to select for mutants that provide resistance. Transposon insertion profiles at the indicated *B. subtilis* genomic regions are shown. Each vertical line indicates an insertion site; its height reflects the number of sequencing reads at this position (maximum height ≥5000). The average number of reads was >40,000. **(A)** The majority of insertions mapped upstream of the *ykoX* gene in an orientation that would increase its transcription. **(B)** Transposon insertions were not enriched upstream of *bcrC* or *uppP* that encode UndPP phosphatases, suggesting that these proteins do not have UndP transport activity, as has been proposed previously (*3, 30*). **(C)** Unlike the Tn-seq screen in a wild-type (*yngC*+) background, in the Δ*yngC* mutant, transposon insertions were not enriched in the genes (*yhdL* and *yhdK*) encoding the anti-σ^M^ factors, consistent with the model that their inactivation provides increased MX2401 resistance by increasing σ^M^-dependent transcription of *yngC*.

**Figure S7.**
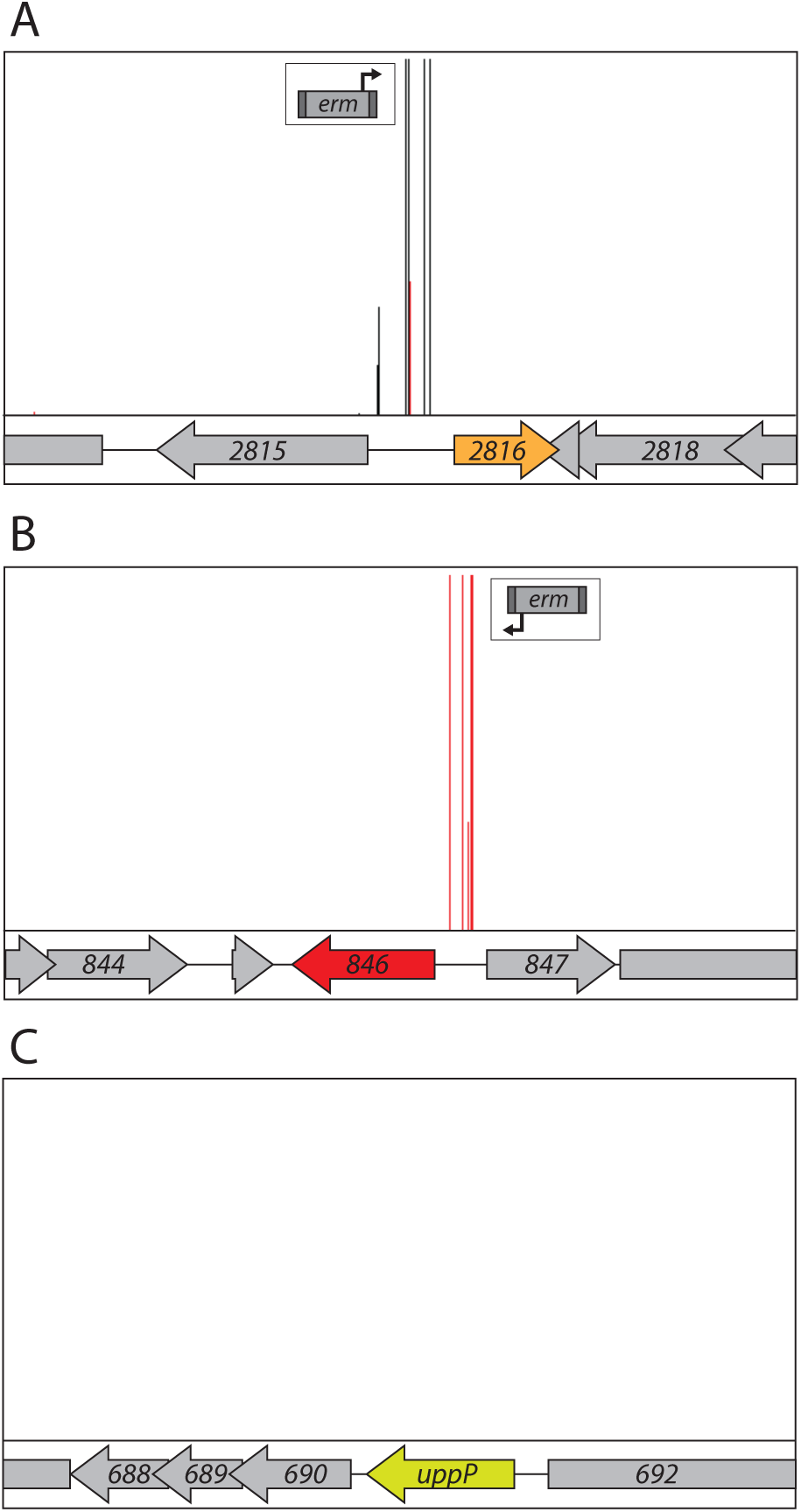
A transposon-sequencing screen identified insertions upstream of *02816* and *00846* that confer resistance to amphomycin in *S. aureus*. A reanalysis of Tn-Seq data from Santiago et al. 2018 (*16*). A library of *S. aureus* mutagenized with a transposon carrying a strong outward facing promoter (P*_tuf_*) was grown in sub-inhibitory concentrations of amphomycin (9.6 µg/mL). Transposon insertion profiles at the indicated *S. aureus* genomic region are shown. Each vertical line indicates an insertion site; its height reflects the number of sequencing reads at this position (maximum height ≥2000). The majority of transposon insertions mapped upstream of *SAOUHSC_02816* (A) and *SAOUHSC_00846* (B) in orientations that are predicted to increase transcription of these two genes. (C) By contrast, insertions were not enriched upstream of the UndPP phosphatase *uppP*, suggesting that this membrane phosphatase does not have UndP transport activity, as has been proposed previously (*3, 30*).

**Figure S8.**
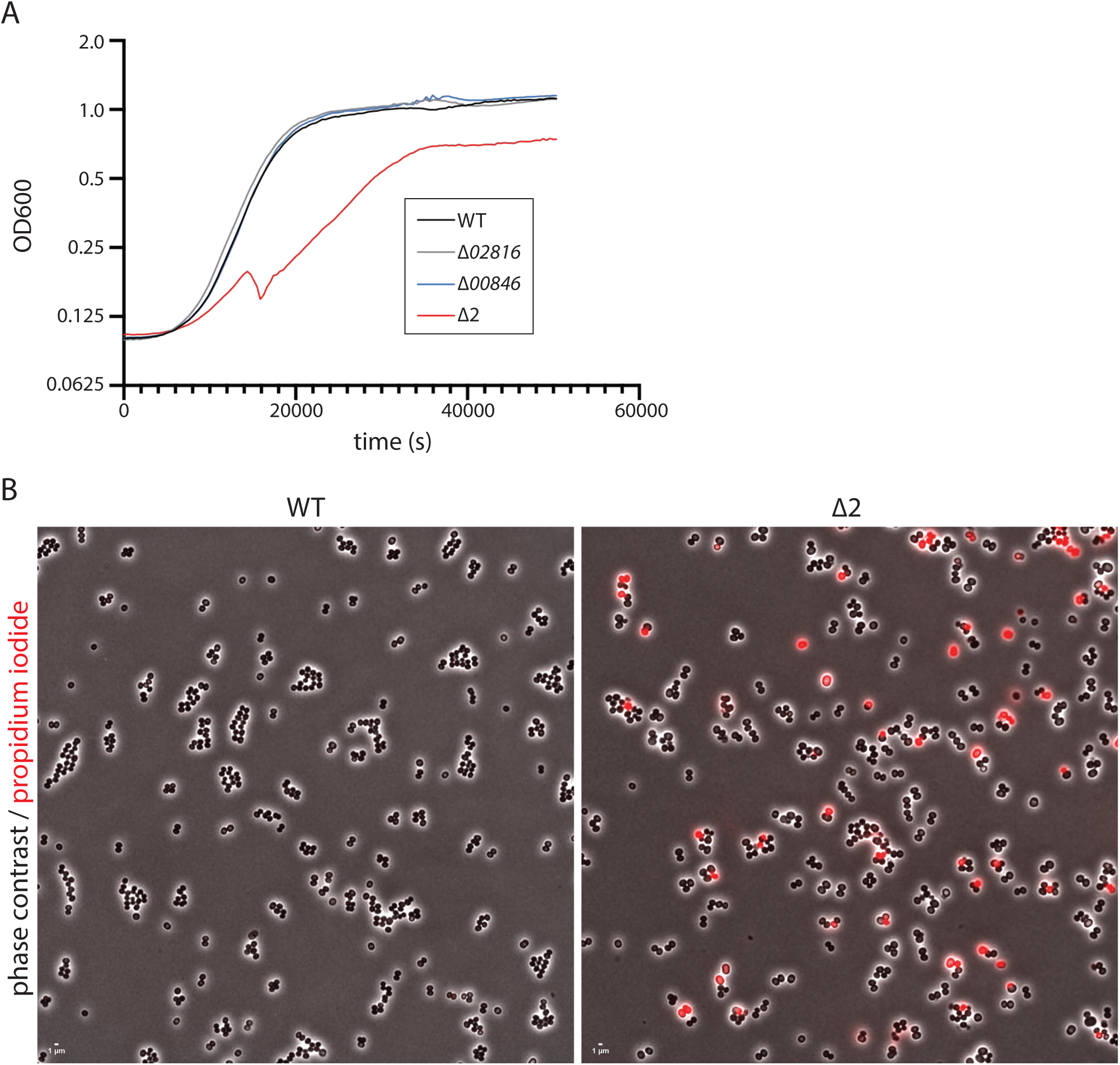
A *ΔpopT ΔuptA S. aureus* mutant has impaired growth and membrane permeability defects. **(A)** Representative growth curves of the indicated strains. Cells were grown in TSB at 37 °C with orbital shaking in a microplate growth reader. **(B)** Representative micrographs of wild-type and the Δ2 (Δ*uptA* Δ*popT*) double mutant. Shown are overlays of phase-contrast and fluorescent images of propidium iodide stained cells. Quantification of the PI-positive cells from several fields of view (>1000 cells per strain) yielded a PI-positive rate of 0.1% for wild-type and 10% for the Δ*uptA* Δ*popT* mutant.

**Figure S9.**
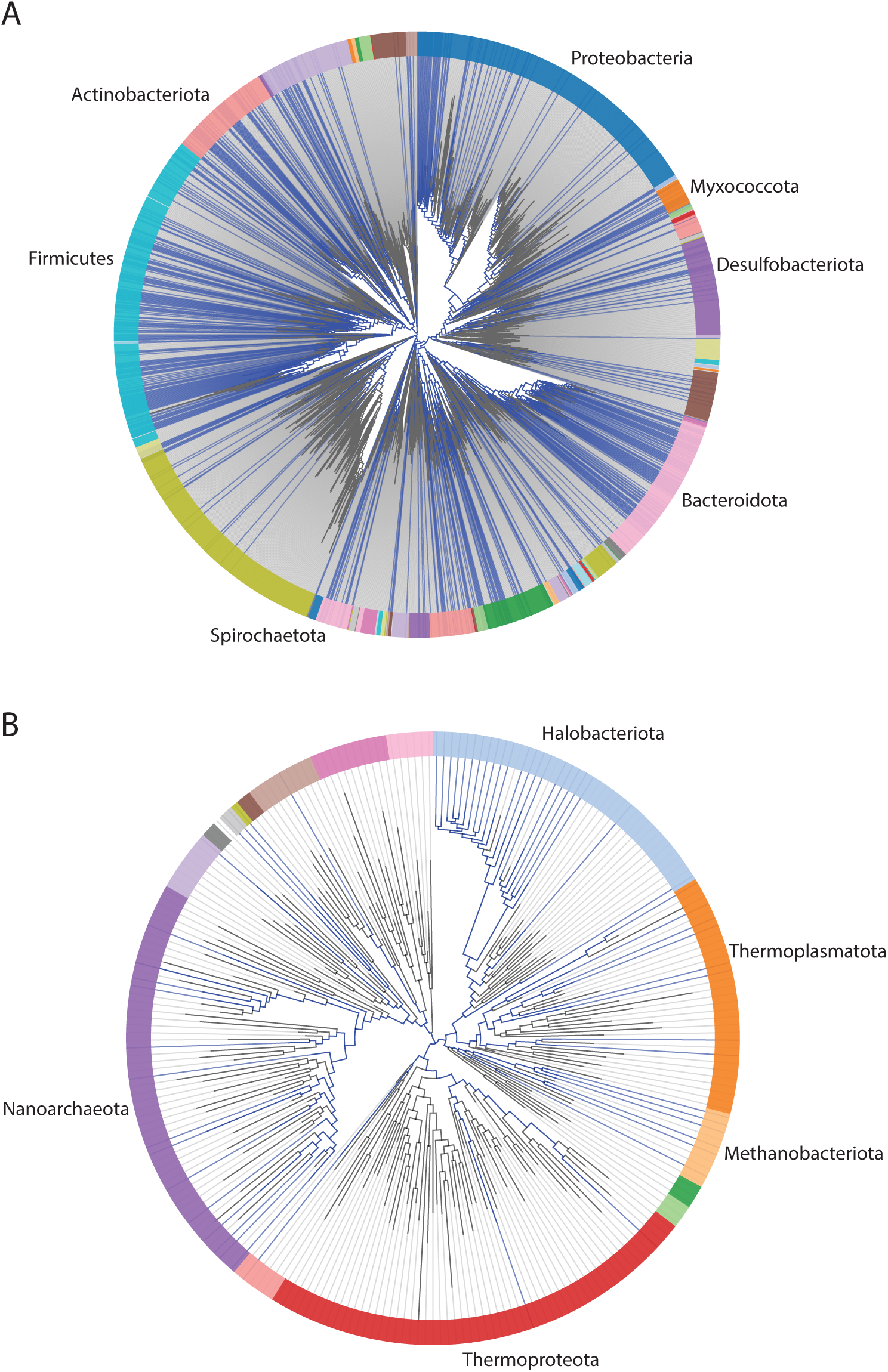
DUF368 is broadly conserved in bacteria and archaea. Dendrograms highlighting the distribution of DUF368 domains in a broad range of bacterial **(A)** and archaeal **(B)** species. All members of the pfam04018 (DUF368) were mapped onto representative phylogenetic trees using AnnoTree (*38*).

**Figure S10.**
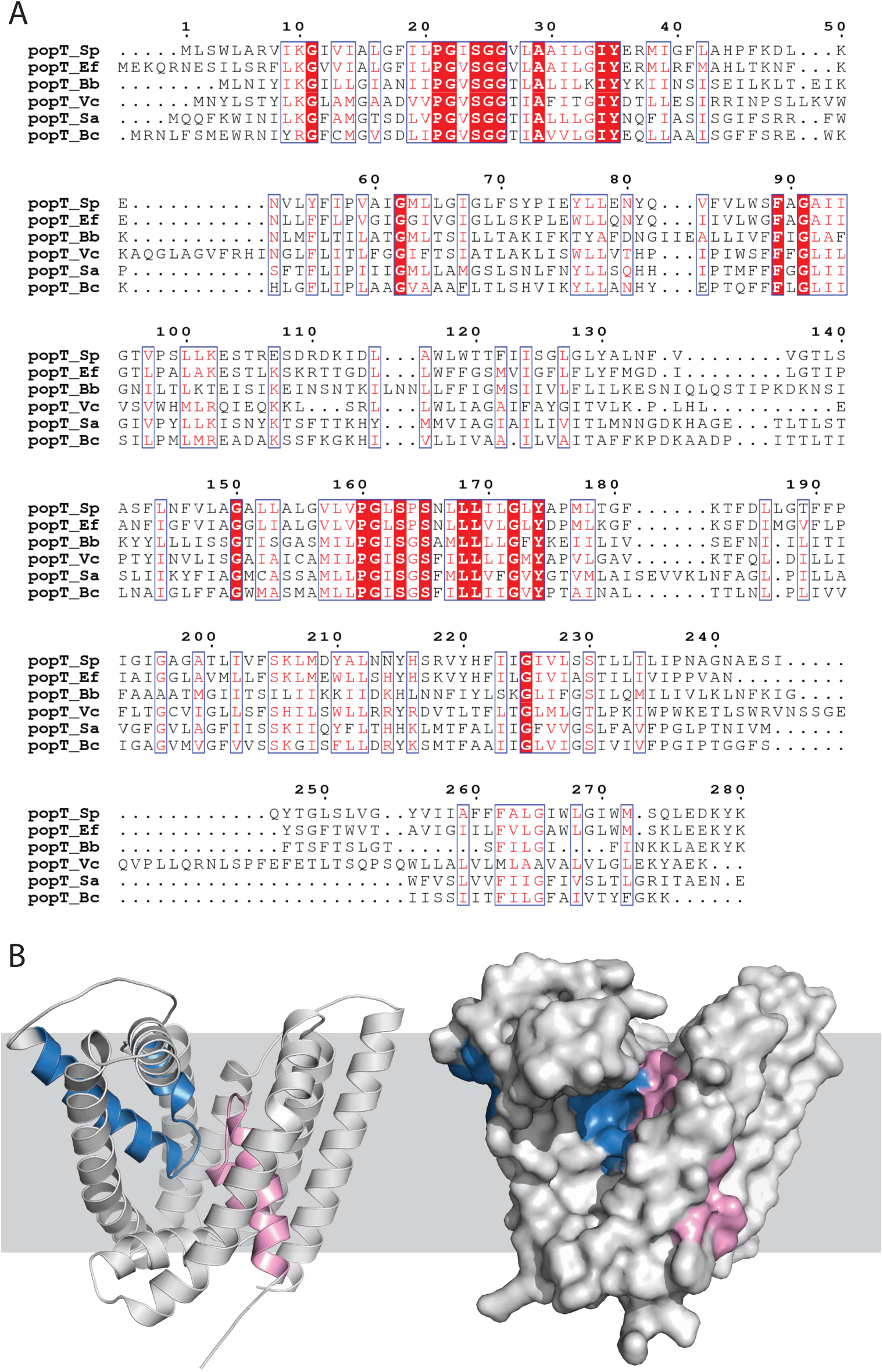
The predicted structure of DUF368 has features of membrane transporters. **(A)** Multiple sequence alignment of DUF368 (PopT) homologs that provide MX2401 resistance. **(B)** Structural models of SAOUHSC_00846 as predicted by AlphaFold2. Highlighted in red and blue are re-entrant helices that are commonly found in membrane embedded transporters (*13*).

**Figure S11.**
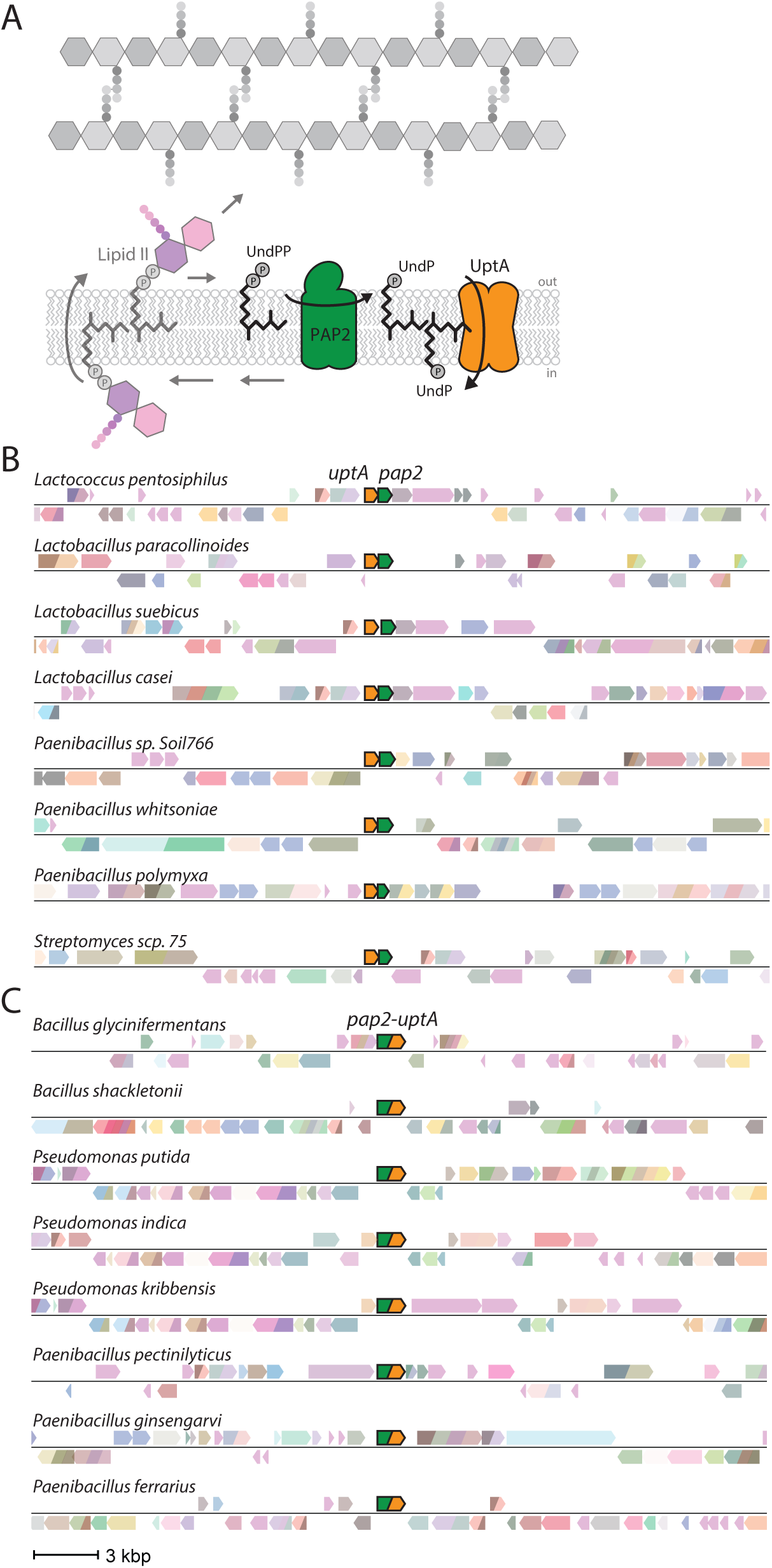
Gene neighborhood analysis reveals that *dedA* transporters are often adjacent to or fused with *pap2* lipid phosphatases. **(A)** Schematic of the lipid II cycle highlighting the role of PAP2 family members like BcrC and DedA family members like YngC (UptA) in dephosphorylating UndPP and flipping UndP across the membrane, respectively. Representative genomic neighborhood analyses showing synteny **(B)** and gene-fusions **(C)** of DedA family members with PAP2 lipid phosphatases. Uniprot IDs for the proteins in the diagram can be found in Table S2.

**Figure S12.**
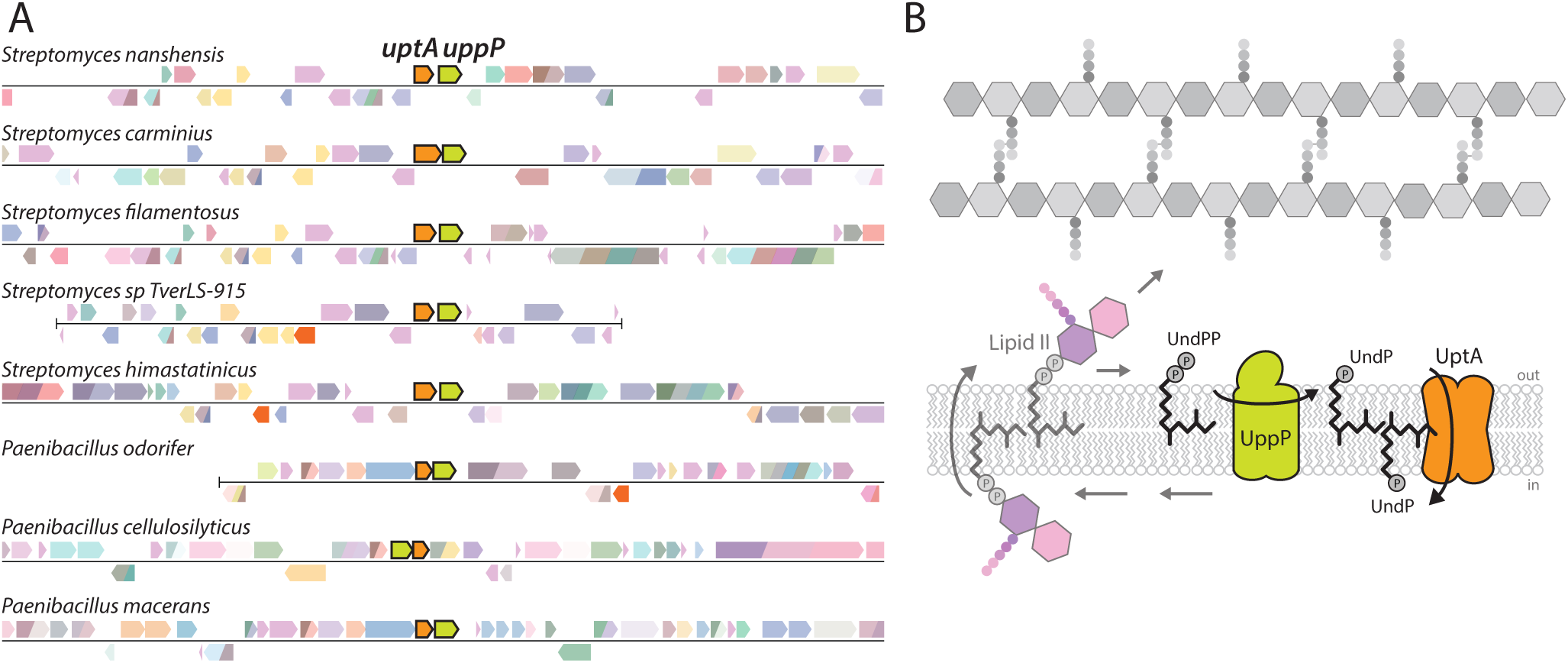
Gene neighborhood analysis reveals that some *dedA* transporters are adjacent to *uppP* undecaprenyl-pyrophosphate phosphatases. (A) Representative genomic neighborhood analysis showing synteny of *dedA* transporters with the undecaprenyl-pyrophosphate phosphatase *uppP*. Most examples of synteny between *dedA* and *uppP* genes are from *Streptomycetes* and *Paenicbacilli* genomes. (B) Schematic of the lipid II cycle highlighting the role of UppP family members and DedA family members like YngC (UptA) in dephosphorylating UndPP and flipping UndP across the membrane, respectively. Uniprot IDs for the proteins included in this diagram can be found in Table S2.

**Figure S13.**
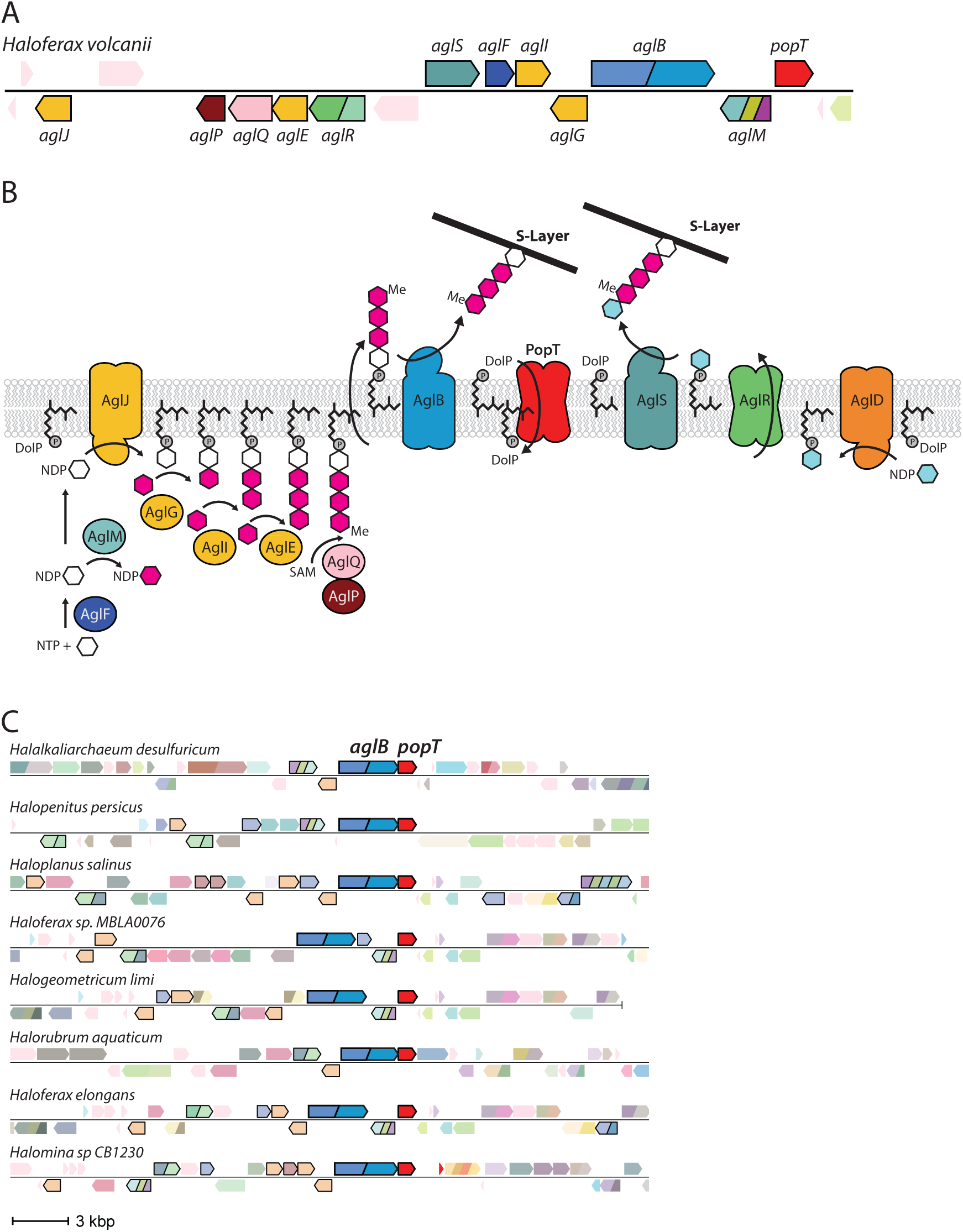
Gene neighborhood analysis reveals *duf368* transporters are often present in archaeal gene clusters involved in S-layer protein glycosylation. (A) Representative genomic neighborhood from *Haloferaz volcanii* with all the characterized genes involved in S-layer protein glycosylation and *popT* highlighted. (B) Schematic of the S-layer protein glycosylation pathway encoded in the *H. volcanii* gene cluster based on *Jarrel et al. 2014* (*24*). PopT is hypothesized to catalyze the recycling of DolP to complete the lipid cycle. (C) Gene neighborhood analysis showing synteny of *duf368* (*popT*) transporters with *aglB* and other genes involved in N-linked protein glycosylation (outlined in black). Uniprot IDs for the proteins included in this diagram can be found in Table S2.

**Figure S14.**
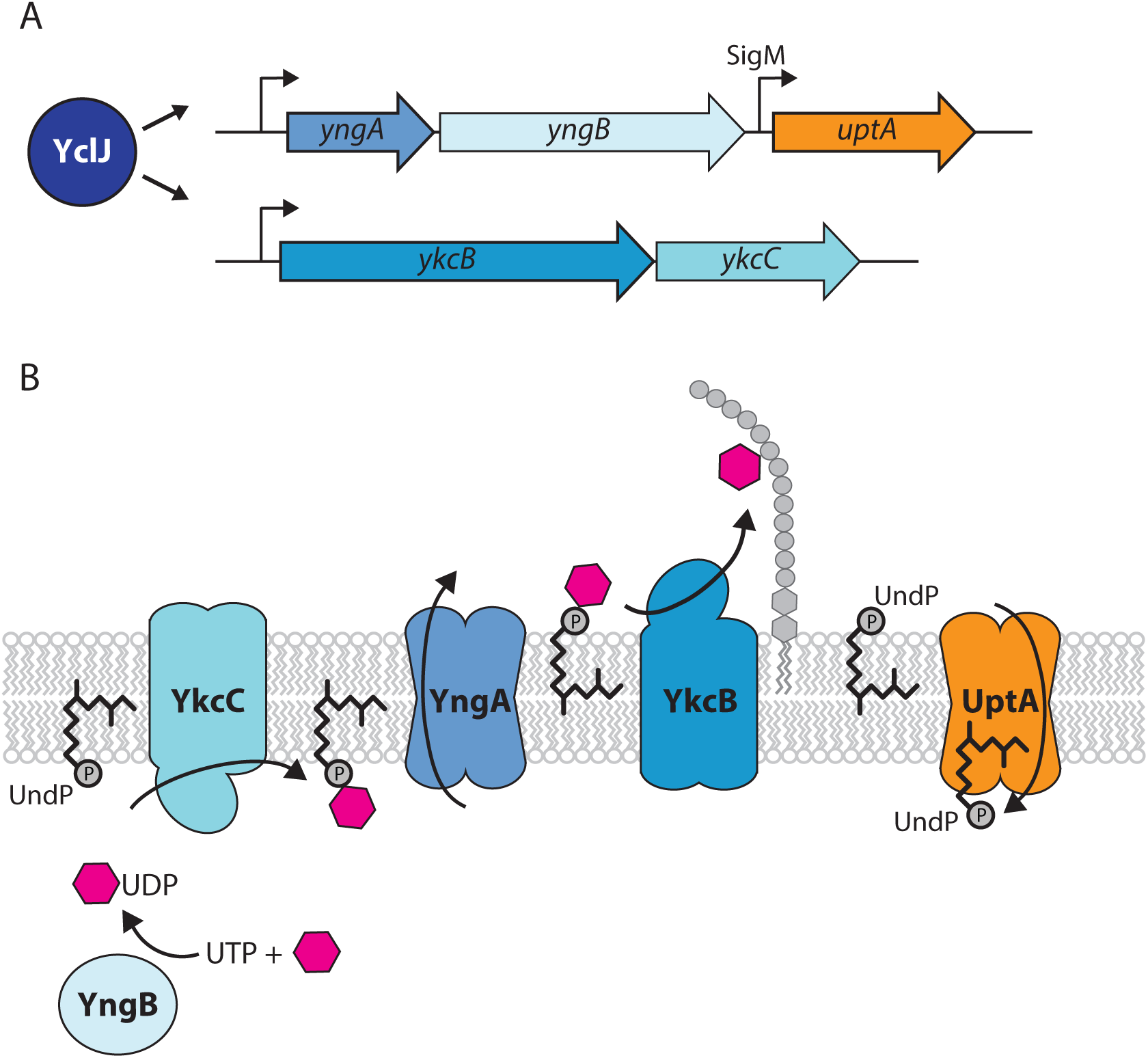
*yngC* is co-regulated with genes encoding a surface polymer glycosylation pathway. **(A)** Cartoon depiction of the regulation of the *yngABC* and *ykcBC* operons by the transcription factor YclJ (*39*). **(B)** Schematic of the putative cell surface glycosylation pathway encoded by YclJ regulon members (*25*). YngA is a member of the GtrA flippase family that transports UndP-linked monosaccharides across lipid bilayers. YngB is a member of the UDP glucose pyrophosphorylase family and has been shown to charge sugars with UDP groups (*25*). YkcB is a member of the glycosyltransferase-39 family that transfers monosaccharides from the UndP carrier to surface polymers. YkcC is a member of the glycosyltransferase-2 family that adds UDP charged monosaccharides onto UndP on the cytoplasmic leaflet of the membrane. The UndP transporter UptA completes the lipid cycle.

**Figure S15.**
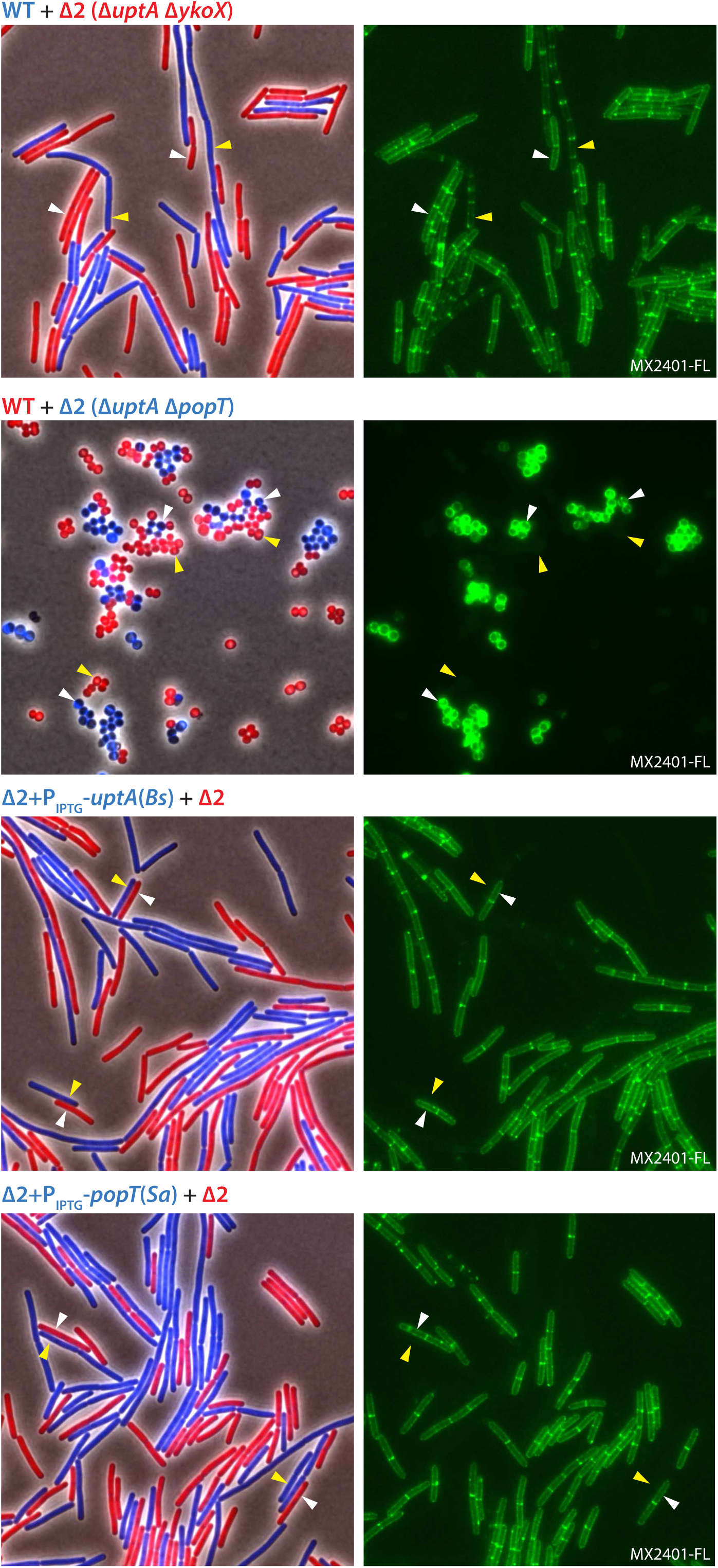
UptA or PopT expression reduces surface exposed UndP in *B. subtilis* and *S. aureus.* Representative microscopy images of the indicated *B. subtilis* or *S. aureus* strains. Two strains expressing different fluorescent proteins (*B. subtilis*) or labeled with different fluorescent D-amino acids (*S. aureus*) were mixed and then stained with fluorescent MX2401 (MX2401-FL). The left panels show overlays of phase contrast and fluorescent images in the red and blue channels to distinguish the two strains. The right panels show MX2401-FL staining. Yellow carets highlight wild-type cells or cells over-expressing UptA(*Bs*) or PopT(*Sa*). White carets highlight cells lacking the UndP transporters.

**Figure S16.**
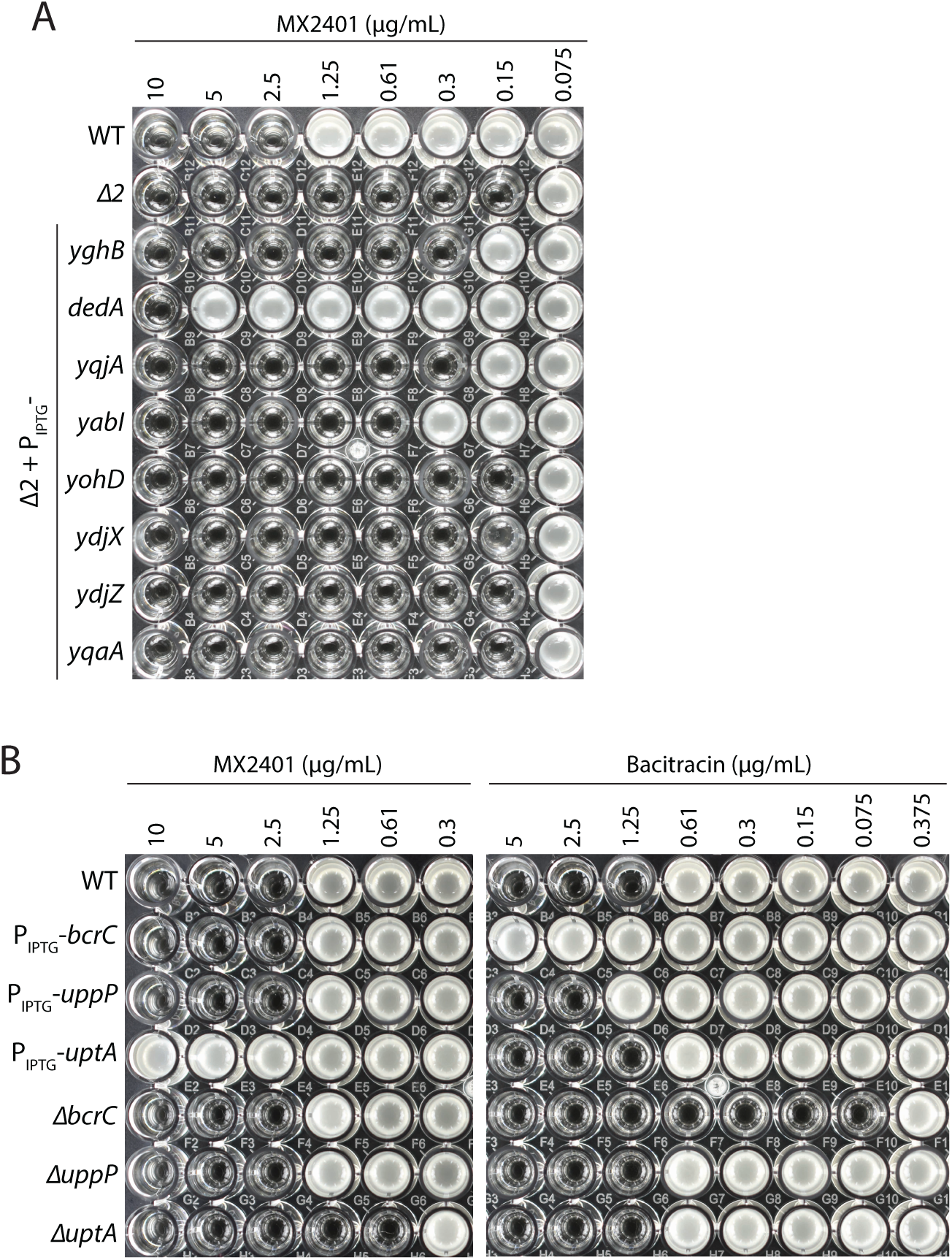
*E. coli* DedA provides MX2401 resistance in *B. subtilis.* **(A)** MIC assays of the indicated strains each expressing one of the eight *E. coli* DedA paralogs in *B. subtilis* lacking *uptA* and *ykoX* (Δ2). Strains were grown in LB with 10 µM IPTG. At 500 µM, cells expressing YqjA and YabI had 4-fold higher MICs (≥8-fold lower than cells expressing DedA). **(B)** Over-expression or deletion of the undecaprenyl-pyrophosphate phosphatase genes *uppP* or *bcrC* in *B. subtilis* do not alter the MIC of MX2401. MIC assays of the indicated *B. subtilis* strains tested with MX2401 and separately with bacitracin that targets UndPP.

**Figure S17.**
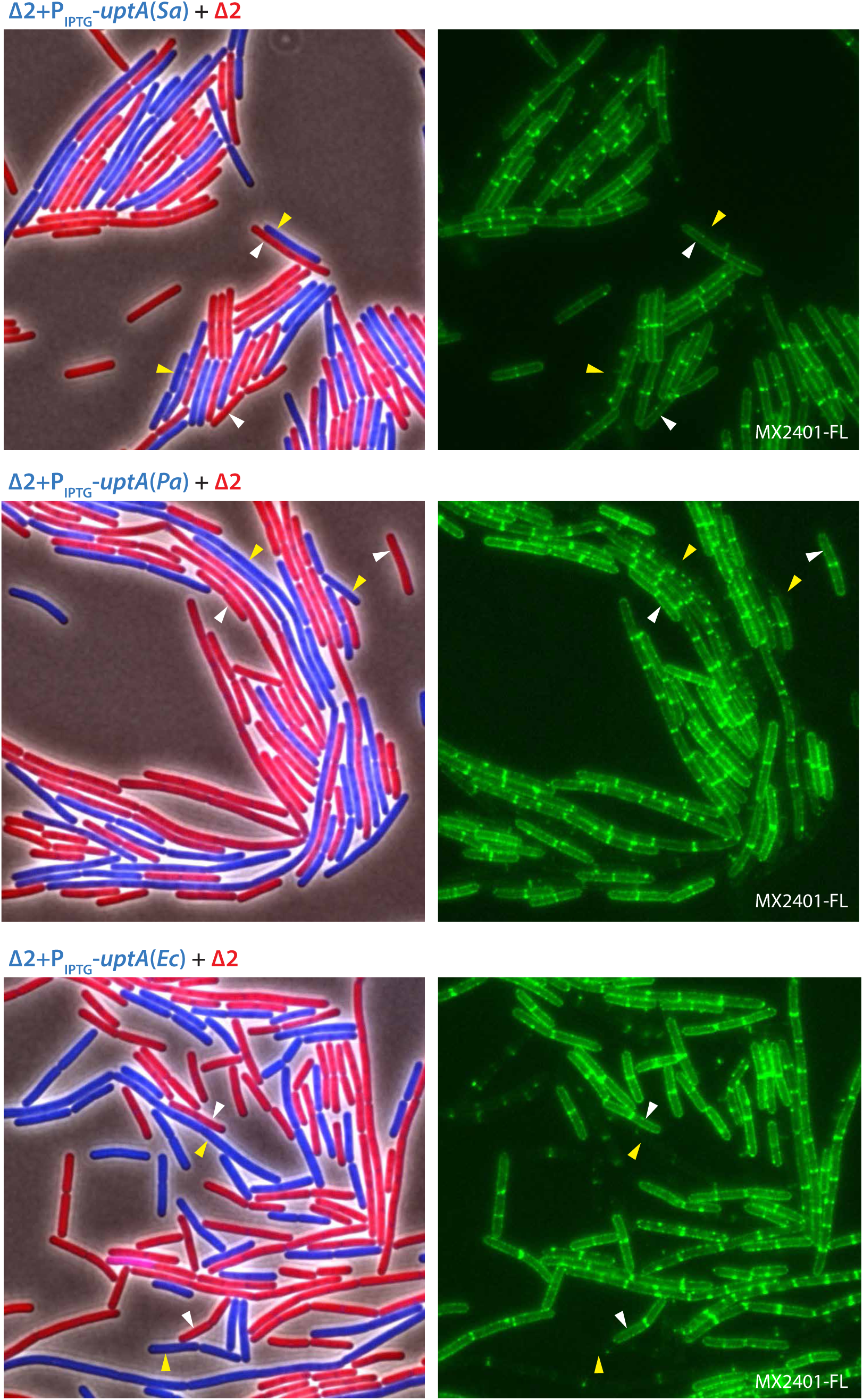
Homologs of UptA from diverse bacterial species can transport UndP in *B. subtilis*. Representative microscopy images of the indicated *B. subtilis* strains. Two strains expressing different fluorescent proteins were mixed and then stained with fluorescent MX2401 (MX2401-FL). The left panels show overlays of phase contrast and fluorescent images in the red and blue channels to distinguish the two strains. The right panels show MX2401-FL staining. Yellow carets highlight cells over-expressing an UptA homolog in a strain lacking *uptA* and *ykoX* (Δ2). White carets highlight cells lacking *uptA* and *ykoX*.

**Figure S18.**
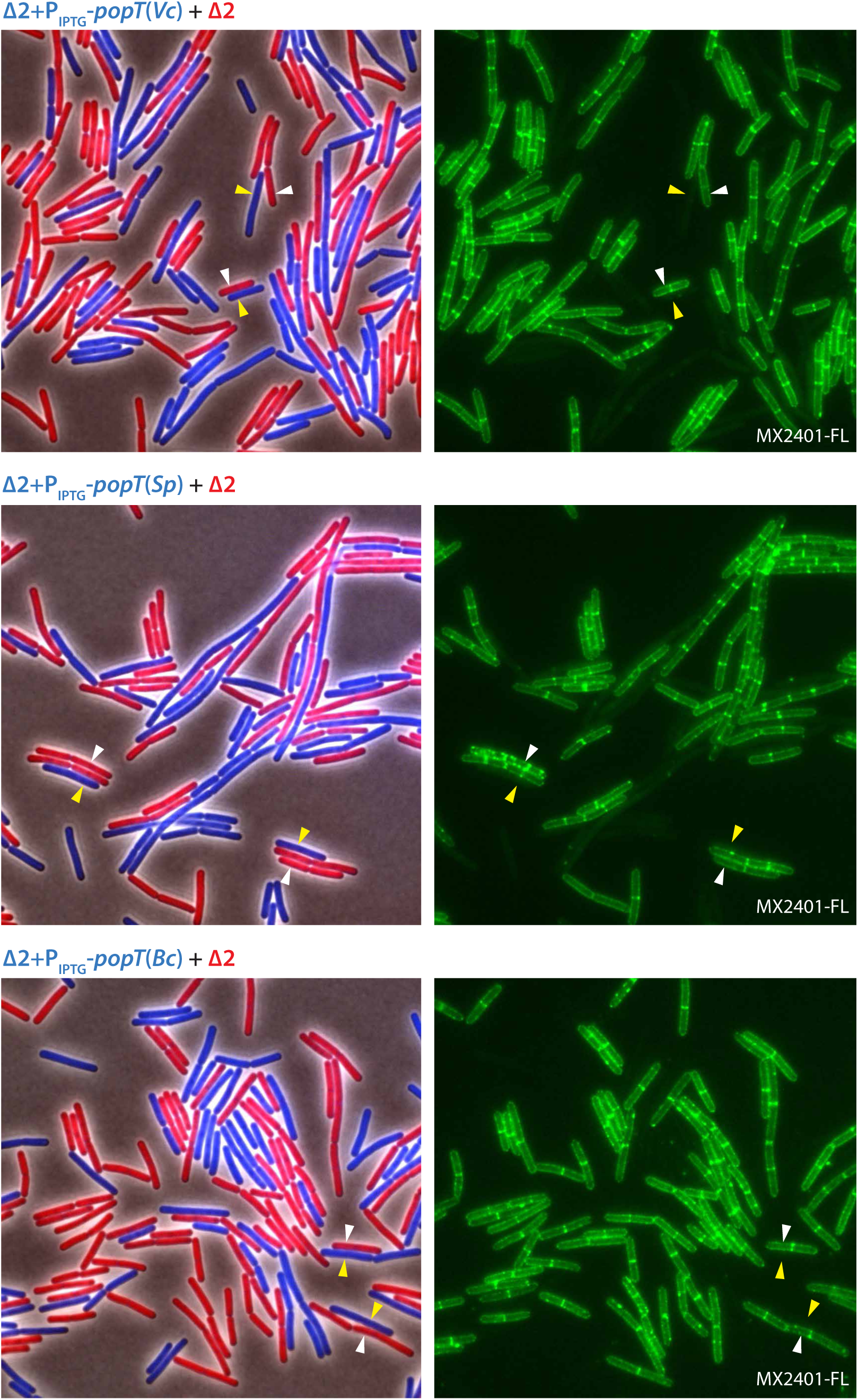
Homologs of PopT from diverse bacterial species can transport UndP in *B. subtilis*. Representative microscopy images of the indicated *B. subtilis* strains. Two strains expressing different fluorescent proteins were mixed and then stained with fluorescent MX2401 (MX2401-FL). The left panels show overlays of phase contrast and fluorescent images in the red and blue channels to distinguish the two strains. The right panels show MX2401-FL staining. Yellow carets highlight cells over-expressing a PopT homolog in a strain lacking *uptA* and *ykoX* (Δ2). White carets highlight cells lacking *uptA* and *ykoX*.

**Table S1A.** Homologies between UptA from B. subtilis and S. aureus and the 8 E. coli DedA paralogs.

| **DedA member that provides MX2401 resistance** | *E.coli* DedA paralogs | E-Value |
| --- | --- | --- |
| YngC (UptA<sup>Bs</sup>)<br>(*Bacillus subtilis* PY79) | DedA (UptA<sup>Ec</sup>) | 9e-13 |
|  | Yabl | 8e-12 |
|  | YohD | 3e-11 |
|  | YqjA | 9e-08 |
|  | YghB | 2e-06 |
|  | YdjZ | ----- |
|  | YdjX | ----- |
|  | YqaA | ----- |
| SAOUHSC_002816 (UptA<sup>Sa</sup>)<br>(*Staphylococcus aureus* NCTC8325) | DedA (UptA<sup>Ec</sup>) | 1e-13 |
|  | YqjA | 3e-12 |
|  | YghB | 1e-08 |
|  | YdjZ | 6e-06 |
|  | Yabl | 3e-06 |
|  | YohD | 1e-06 |
|  | YqaA | 0.002 |
|  | YdjX | ----- |

**Table S1B.**
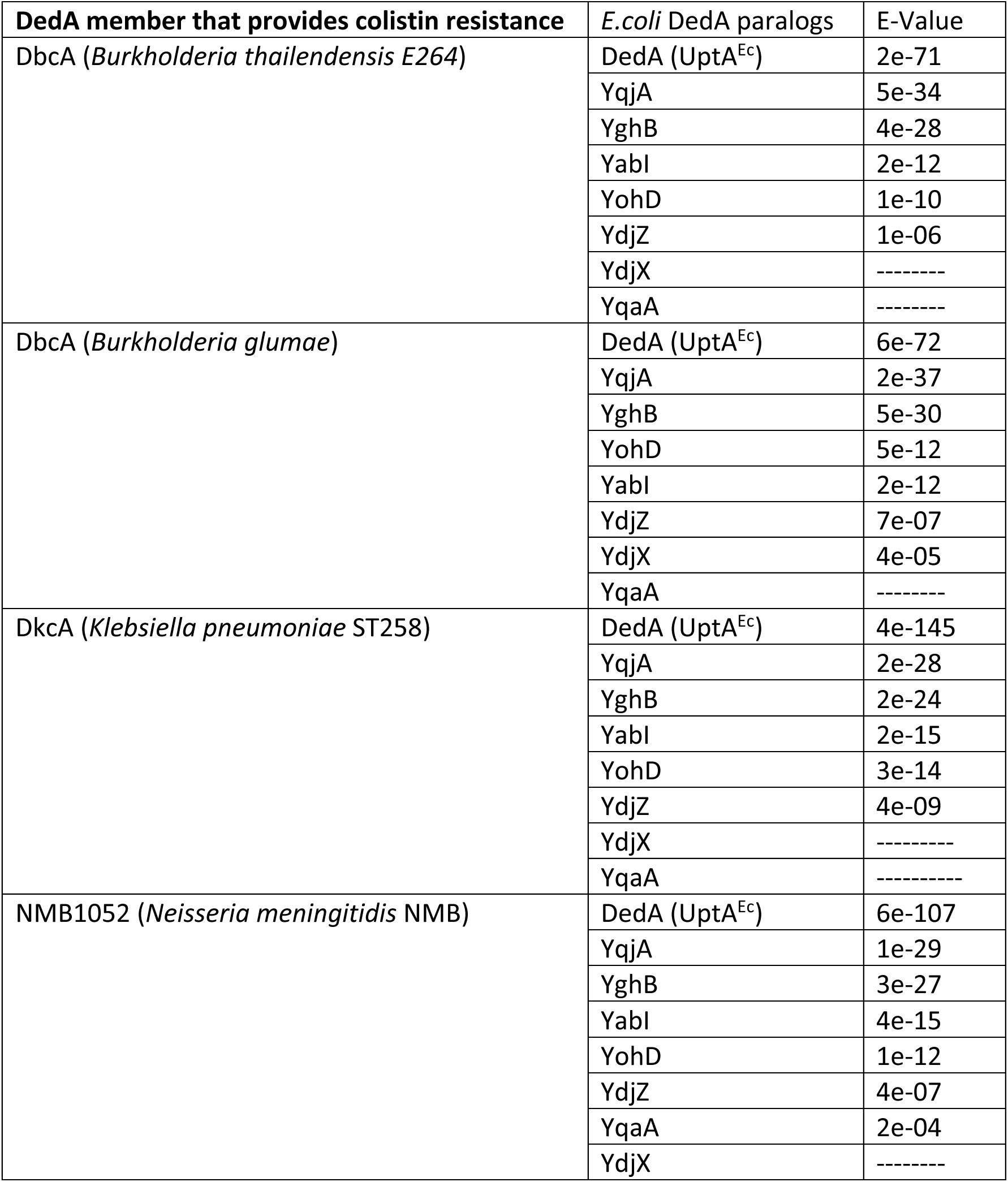
Homologies between the DedA family members in B. thailendensis, B. glumae, and K. pneumoniae and the 8 E. coli DedA paralogs.

**Table S2.**
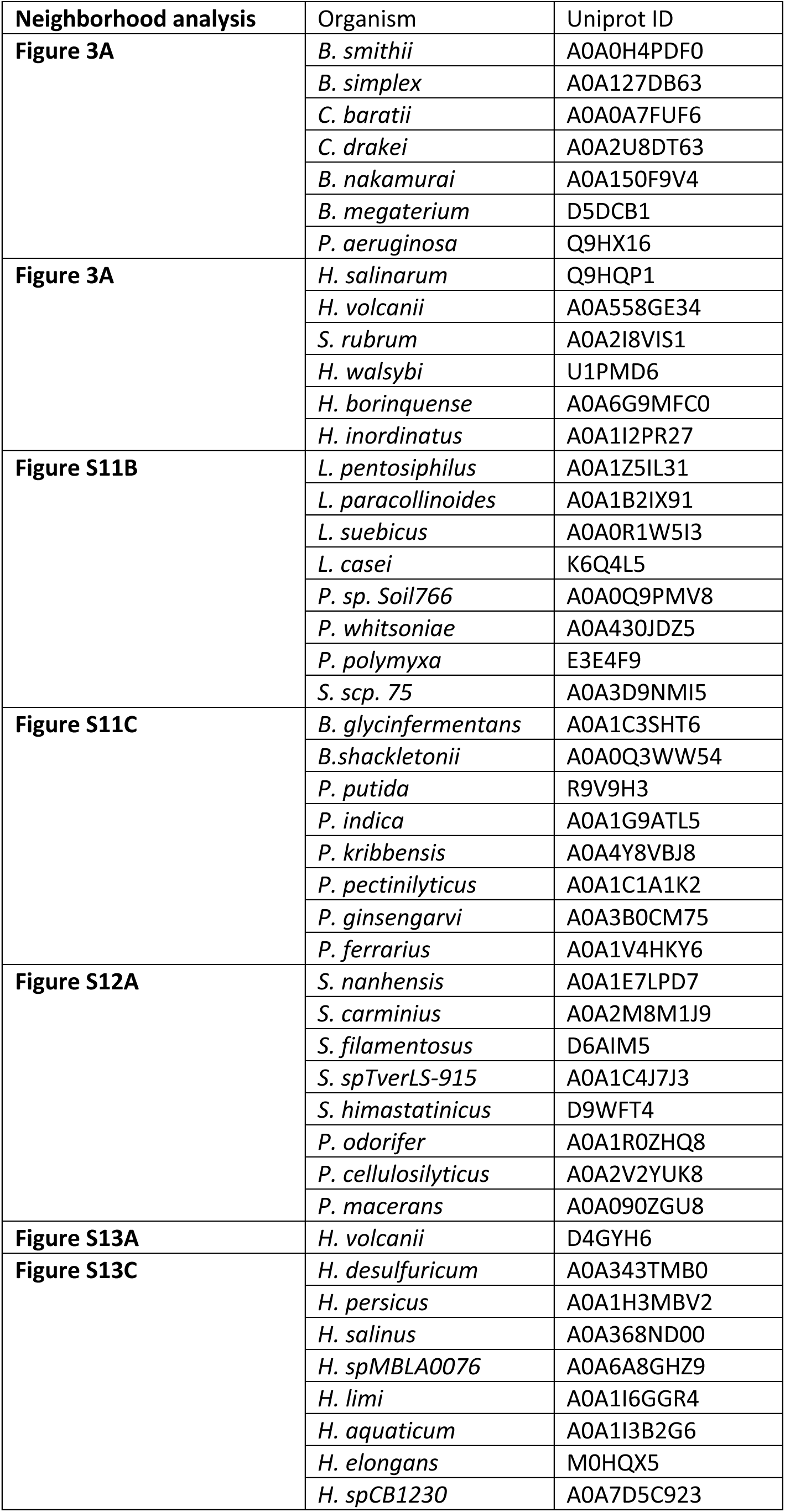
Uniprot IDs for the proteins included in gene neighborhood analyses.

**Table S3.** Strains used in this study.

| Strain | Background | Genotype | Source | Figures |
| --- | --- | --- | --- | --- |
| ATP001 | *S. aureus*<br>RN4220 | wild-type | (1) | 2ce, 4a, S8ab |
| BIR003 | *B. subtilis*<br>PY79 | wild-type | (2) | S3e |
| BIR019 | *B. subtilis*<br>PY79 | $\Delta(\text{sigM-yhdL-yhdK})::\text{erm}$ | This study | S3e |
| BIR0334 | *B. subtilis*<br>PY79 | *sacA::Pveg-mTagBFP (phleo), amyE::Pamj-yfp (cat)* | This study | 2bf, 4b, S2c, S4b, S10c, S16 |
| BIR0627 | *B. subtilis*<br>PY79 | *sacA::Pveg-mTagBFP (phleo), amyE::Pamj-yfp (cat), uppP::erm* | This study | S16 |
| BIR0634 | *B. subtilis*<br>PY79 | *sacA::Pveg-mTagBFP (phleo), amyE::Pamj-yfp (cat), ycgO::Pspank-uppS (spec), uppS::tet* | This study | S4b |
| BIR0644 | *B. subtilis*<br>PY79 | *sacA::Pveg-mTagBFP (phleo), amyE::Pamj-yfp (cat), yngC::erm* | This study | 2b, S3e, S4b, S16b |
| BIR0645 | *B. subtilis*<br>PY79 | *sacA::Pveg-mTagBFP (phleo), amyE::Pamj-yfp (cat), ykoX::kan* | This study | 2b |
| BIR0646 | *B. subtilis*<br>PY79 | *sacA::Pveg-mTagBFP (phleo), amyE::Pamj-yfp (cat), ybfM::tet* | This study |  |
| BIR0648 | *B. subtilis*<br>PY79 | *sacA::Pveg-mTagBFP (phleo), amyE::Pamj-yfp (cat), yngC::erm, ykoX::kan* | This study | 2bf, 4b, S2c, S10c, S16a |
| BIR0650 | *B. subtilis*<br>PY79 | *sacA::Pveg-mTagBFP (phleo), amyE::Pamj-yfp (cat), ycgO::Phyperspank-yngC (spec)* | This study | 2b |
| BIR0668 | *B. subtilis*<br>PY79 | *sacA::Pveg-mTagBFP (phleo), amyE::Pamj-yfp (cat), bcrC::erm* | This study | S16b |
| BIR0672 | *B. subtilis*<br>PY79 | *sacA::Pveg-mTagBFP (phleo), amyE::Pamj-yfp (cat), yngC::tet* | This study | S6 |
| BIR0683 | *S. aureus*<br>RN4220 | $\Delta 02816::\text{spec}$ | This study | 2cd, S8a |
| BIR0688 | *S. aureus*<br>RN4220 | $\Delta 00846::\text{kan}$ | This study | 2cd, S8a |
| BIR0691 | *S. aureus*<br>RN4220 | *attB::Ptet-02816 (cat)* | This study | 2c |
| BIR0695 | *S. aureus*<br>RN4220 | *attB::Ptet-00846 (cat)* | This study | 2c |
| BIR0712 | *B. subtilis*<br>PY79 | *sacA::Pveg-mTagBFP (phleo), amyE::Pamj-yfp (cat), yngC::lox72, ykoX::lox72, ybfM::lox72, yhjE::kan, yqeD::erm, ytxB::tet* | This study | 2b |
| BIR0715 | *S. aureus*<br>RN4220 | $\Delta 02816::\text{spec}, \Delta 00846::\text{kan}$ | This study | 2cde, 4a, S8ab |
| BIR0769 | *B. subtilis*<br>PY79 | *sacA::Pveg-mTagBFP (phleo), amyE::Pamj-yfp (cat), bceAB::kan* | This study | S16b |
| BIR0852 | *B. subtilis*<br>PY79 | *sacA::Pveg-mTagBFP (phleo), amyE::Pamj-yfp (cat), ycgO::Pspank-ispH (spec), ispH::kan* | This study | S4b |
| BIR1113 | *B. subtilis*<br>PY79 | *sacA::Pveg-mTagBFP (phleo), amyE::Pamj-yfp (cat), yngC::erm, ykoX::kan, ycgO::Phyperspank-yngC (spec)* | This study | 2f, 4b, S2c |
| BIR1114 | *B. subtilis*<br>PY79 | *sacA::Pveg-mTagBFP (phleo), amyE::Pamj-yfp (cat), yngC::erm, ykoX::kan, ycgO::Phyperspank-ykoX (spec)* | This study | 2f |
| BIR1116 | *B. subtilis*<br>PY79 | *sacA::Pveg-mTagBFP (phleo), amyE::Pamj-yfp (cat), yngC::erm, ykoX::kan, ycgO::Phyperspank-02816 (spec)* | This study | 2f |
| BIR1117 | *B. subtilis*<br>PY79 | *sacA::Pveg-mTagBFP (phleo), amyE::Pamj-yfp (cat), yngC::erm, ykoX::kan, ycgO::Phyperspank-00846 (spec)* | This study | 2f, S10C |
| BIR1118 | *B. subtilis*<br>PY79 | *sacA::Pveg-mTagBFP (phleo), amyE::Pamj-yfp (cat), yngC::erm, ykoX::kan, ycgO::Phyperspank- yghB(Ec) (spec)* | This study | S16a |
| BIR1119 | *B. subtilis*<br>PY79 | *sacA::Pveg-mTagBFP (phleo), amyE::Pamj-yfp (cat), yngC::erm, ykoX::kan, ycgO::Phyperspank- yqjA(Ec) (spec)* | This study | S16a |
| BIR1120 | *B. subtilis*<br>PY79 | *sacA::Pveg-mTagBFP (phleo), amyE::Pamj-yfp (cat), yngC::erm, ykoX::kan, ycgO::Phyperspank-yngC-his10 (spec)* | This study | S2cd |
| BIR1132 | *B. subtilis*<br>PY79 | *sacA::Pveg-mTagBFP (phleo), amyE::Pamj-yfp (cat), yngC::erm, ykoX::kan, ycgO::Phyperspank-yngC(R112A)-his10 (spec)* | This study | S2cd |
| BIR1133 | *B. subtilis*<br>PY79 | *sacA::Pveg-mTagBFP (phleo), amyE::Pamj-yfp (cat), yngC::erm, ykoX::kan, ycgO::Phyperspank-yngC(R118A)-his10 (spec)* | This study | S2cd |
| BIR1134 | *B. subtilis*<br>PY79 | *sacA::Pveg-mTagBFP (phleo), amyE::Pamj-yfp (cat), yngC::erm, ykoX::kan, ycgO::Phyperspank-yngC(R112A,R118A)-his10 (spec)* | This study | S2cd |
| BIR1186 | *B. subtilis*<br>PY79 | *sacA::Pveg-mTagBFP (phleo), yngC::erm, ykoX::kan, ycgO::Phyperspank-yngC(spec)* | This study | 4a, S15 |
| BIR1187 | *B. subtilis*<br>PY79 | *sacA::Pveg-mTagBFP (phleo), yngC::erm, ykoX::kan, ycgO::Phyperspank-00846 (spec)* | This study | 4a, S15 |
| BIR1191 | *B. subtilis*<br>PY79 | *sacA::Pveg-mTagBFP (phleo), yngC::erm, ykoX::kan, ycgO::Phyperspank-02816 (spec)* | This study | S17 |
| BIR1193 | *B. subtilis*<br>PY79 | *sacA::Pveg-mCherry (phleo), ykoX::kan, yngC::erm* | This study | 4a, S15, S17, S18 |
| BIR1218 | *B. subtilis*<br>PY79 | *sacA::Pveg-mTagBFP (phleo), amyE::Pamj-yfp (cat), yngC::erm, ykoX::kan, ycgO::Phyper-uptA(A.baumannii) (spec)* | This study | 4b |
| BIR1219 | *B. subtilis*<br>PY79 | *sacA::Pveg-mTagBFP (phleo), yngC::erm, ykoX::kan, ycgO::Phyperspank-popT(B.burgdorferi) (spec)* | This study | 4c |
| BIR1227 | *B. subtilis*<br>PY79 | *sacA::Pveg-mTagBFP (phleo), yngC::erm, ykoX::kan, ycgO::Phyperspank-popT(C.glutamicum) (spec)* | This study | 4c |
| BIR1232 | *B. subtilis*<br>PY79 | *amyE::PyngC-optRBS-(ATG)-yfp (cat), sacA::Pveg-mTag-BFP(phleo)* | This study | S3bde |
| BIR1233 | *B. subtilis*<br>PY79 | *amyE::PyngC-optRBS-(ATG)-yfp (cat), Δmlk::erm, sacA::Pveg-mTag-BFP(phleo)* | This study | S3cde |
| BIR1252 | *B. subtilis*<br>PY79 | *sacA::Pveg-mTagBFP (phleo), amyE::Pamj-yfp (cat), ycgO::Pspank-uppS(spec), uppS::tet, yngC::kan* | This study | S4b |
| BIR1259 | *B. subtilis*<br>PY79 | *sacA::Pveg-mTagBFP (phleo), amyE::Pamj-yfp (cat), ycgO::Pspank-ispH (spec), ispH::kan, yngC::tet* | This study | S4b |
| BIR1267 | *B. subtilis*<br>PY79 | *sacA::Pveg-mTagBFP (phleo), amyE::Pamj-yfp (cat), yngC::erm, ykoX::kan, ycgO::Phyperspank-dedA(E.coli) (spec)* | This study | S16a |
| BIR1269 | *B. subtilis*<br>PY79 | *sacA::Pveg-mTagBFP (phleo), amyE::Pamj-yfp (cat), yngC::erm, ykoX::kan, ycgO::Phyperspank-uptA(B.cereus) (spec)* | This study | 4b |
| BIR1270 | *B. subtilis*<br>PY79 | *sacA::Pveg-mTagBFP (phleo), yngC::erm, ykoX::kan, ycgO::Phyperspank-popT(E.faecium) (spec)* | This study | 4c |
| BIR1273 | *B. subtilis*<br>PY79 | *sacA::Pveg-mTagBFP (phleo), amyE::Pamj-yfp (cat), yngC::erm, ykoX::kan, ycgO::Phyperspank-yabl(E.coli) (spec)* | This study | S16a |
| BIR1274 | *B. subtilis*<br>PY79 | *sacA::Pveg-mTagBFP (phleo), amyE::Pamj-yfp (cat), yngC::erm, ykoX::kan, ycgO::Phyperspank-yohD(E.coli) (spec)* | This study | S16a |
| BIR1275 | *B. subtilis*<br>PY79 | *sacA::Pveg-mTagBFP (phleo), amyE::Pamj-yfp (cat), yngC::erm, ykoX::kan, ycgO::Phyperspank-ydjX(E.coli) (spec)* | This study | S16a |
| BIR1276 | *B. subtilis*<br>PY79 | *sacA::Pveg-mTagBFP (phleo), amyE::Pamj-yfp (cat), yngC::erm, ykoX::kan, ycgO::Phyperspank-ydjZ(E.coli) (spec)* | This study | S16a |
| BIR1277 | *B. subtilis*<br>PY79 | *sacA::Pveg-mTagBFP (phleo), amyE::Pamj-yfp (cat), yngC::erm, ykoX::kan, ycgO::Phyperspank-yqaA(E.coli) (spec)* | This study | S16a |
| BDR2660 | *B. subtilis*<br>PY79 | *sacA::Pveg-mTagBFP (phleo)* | Unpublished | 4a, S15 |
| BIR1279 | *S. aureus*<br>RN4220 | *attB::Ptet-02816 (cat), Δ02816::spec, Δ00846::kan* | This study | 2cd |
| BIR1280 | *S. aureus*<br>RN4220 | *attB::Ptet-00846 (cat), Δ02816::spec, Δ00846::kan* | This study | 2cd |
| BIR1298 | *B. subtilis*<br>PY79 | *sacA::Pveg-mTagBFP (phleo), amyE::Pamj-yfp (cat), ycgO::Phyperspank-bcrC (spec)* | This study | S16b |
| BIR1299 | *B. subtilis*<br>PY79 | *sacA::Pveg-mTagBFP (phleo), amyE::Pamj-yfp (cat), ycgO::Phyperspank-uppP (spec)* | This study | S16b |
| BIR1305 | *B. subtilis*<br>PY79 | *sacA::Pveg-mTagBFP (phleo), yngC::erm, ykoX::kan, ycgO::Phyperspank-popT(S.pneumoniae) (spec)* | This study | 4c, S18 |
| BIR1306 | *B. subtilis*<br>PY79 | *sacA::Pveg-mTagBFP (phleo), yngC::erm, ykoX::kan, ycgO::Phyperspank-popT(V.cholerae) (spec)* | This study | 4c, S18 |
| BIR1307 | *B. subtilis*<br>PY79 | *sacA::Pveg-mTagBFP (phleo), yngC::erm, ykoX::kan, ycgO::Phyperspank-popT(B.cereus) (spec)* | This study | 4c, S18 |
| BIR1308 | *B. subtilis*<br>PY79 | *sacA::Pveg-mTagBFP (phleo), yngC::erm, ykoX::kan, ycgO::Phyperspank -uptA(P.aeruginosa) (spec)* | This study | 4b, S17 |
| BIR1309 | *B. subtilis*<br>PY79 | *sacA::Pveg-mTagBFP (phleo), yngC::erm, ykoX::kan, ycgO::Phyperspank-dedA(E.coli) (spec)* | This study | 4b, S17 |
| BIR1310 | *B. subtilis*<br>PY79 | *sacA::Pveg-mTagBFP (phleo), amyE::Pamj-yfp (cat), yqeD::erm* | This study | S5 |
| BIR1311 | *B. subtilis*<br>PY79 | *sacA::Pveg-mTagBFP (phleo), amyE::Pamj-yfp (cat), yhjE::kan* | This study | S5 |
| BIR1312 | *B. subtilis*<br>PY79 | *sacA::Pveg-mTagBFP (phleo), amyE::Pamj-yfp (cat), ytxB::tet* | This study | S5 |

1. T. Pang, X. Wang, H. C. Lim, T. G. Bernhardt, D. Z. Rudner, The nucleoid occlusion factor Noc controls DNA replication initiation in *Staphylococcus aureus*. *Plos Genet.* 13, e1006908 (2017).
2. P. Youngman, J. B. Perkins, R. Losick, Construction of a cloning site near one end of Tn917 into which foreign DNA may be inserted without affecting transposition in *Bacillus subtilis* or expression of the transposon-borne *erm* gene. *Plasmid.* 12, 1–9 (1984).

**Table S4.** Plasmids used in this study.

| Plasmid | Description | Source |
| --- | --- | --- |
| pAM155 | amyE::Pamj-yfp(cat)(amp) | (1) |
| pDR244 | Ppa-cre, ori(ts) (spec) (amp) | (2) |
| pIR211 | ycgO::Pspank-uppS(spec)(amp) | This study |
| pIR224 | ycgO::Phyperspank-yngC (spec) (amp) | This study |
| pIR226 | ycgO::Phyperspank-ykoX (spec) (amp) | This study |
| pIR233 | ycgO::Phyperspank-bcrC (spec) (amp) | This study |
| pIR234 | ycgO::Phyperspank-uppP (spec) (amp) | This study |
| pIR235 | ycgO::Phyperspank-yghB(spec)(amp) | This study |
| pIR236 | ycgO::Phyperspank-yqjA(spec)(amp) | This study |
| pIR237 | Pmad3.1-2816::spec (erm)(amp) | This study |
| pIR238 | Pmad3.1-846::kan (erm)(amp) | This study |
| pIR239 | attB::Ptet-02816 (cat) (amp) | This study |
| pIR240 | attB::Ptet-00846 (cat) (amp) | This study |
| pIR242 | Himar1C9 IR-specPpen-IR terminators (erm)(amp) | (3) |
| pIR278 | ycgO::Pspank-ispH (spec)(amp) | This study |
| pIR352 | ycgO::Phyperspank-yngC-his10 (spec)(amp) | This study |
| pIR361 | Pmad3.1-PyngC(ΔPsigM) (erm)(amp) | This study |
| pIR385 | amyE::PyngC-yfp (cat)(amp) | This study |
| pIR388 | ycgO::Phyperspank-popT(Vc) (spec)(amp) | This study |
| pIR392 | ycgO::Phyperspank-uptA(Ab)(spec)(amp) | This study |
| pIR393 | ycgO::Phyperspank-popT(Bb) (spec)(amp) | This study |
| pIR397 | ycgO::Phyperspank-uptA(Pa) (spec)(amp) | This study |
| pIR399 | ycgO::Phyperspank-popT(Bc) (spec)(amp) | This study |
| pIR400 | ycgO::Phyperspank-popT(Sp)(spec)(amp) | This study |
| pIR401 | ycgO::Phyperspank-popT(Cg) (spec)(amp) | This study |
| pIR403 | ycgO::Phyperspank-dedA(Ec) (spec)(amp) | This study |
| pIR405 | ycgO::Phyperspank-uptA(Bc) (spec)(amp) | This study |
| pIR406 | ycgO::Phyperspank-yabl(Ec) (spec)(amp) | This study |
| pIR407 | ycgO::Phyperspank-yohD(Ec) (spec)(amp) | This study |
| pIR408 | ycgO::Phyperspank-ydjX(Ec) (spec)(amp) | This study |
| pIR409 | ycgO::Phyperspank-ydjZ(Ec) (spec)(amp) | This study |
| pIR410 | ycgO::Phyperspank-yqaA(Ec) (spec)(amp) | This study |
| pIR411 | ycgO::Phyperspank-popT(Ef) (spec)(amp) | This study |

1. A. J. Meeske, L.-T. Sham, H. Kimsey, B.-M. Koo, C. A. Gross, T. G. Bernhardt, D. Z. Rudner, MurJ and a novel lipid II flippase are required for cell wall biogenesis in *Bacillus subtilis*. *Proc National Acad Sci.* 112, 6437–6442 (2015). 2. X. Wang, T. B. K. Le, B. R. Lajoie, J. Dekker, M. T. Laub, D. Z. Rudner, Condensin promotes the juxtaposition of DNA flanking its loading site in *Bacillus subtilis*. *Gene Dev.* 29, 1661–1675 (2015). 3. G. S. Dobihal, J. Flores-Kim, I. J. Roney, X. Wang, D. Z. Rudner, The WalR-WalK signaling pathway modulates the activities of both CwlO and LytE through control of the peptidoglycan deacetylase PdaC in *Bacillus subtilis*. *J Bacteriol*, JB0053321 (2021).

**Table S5.**
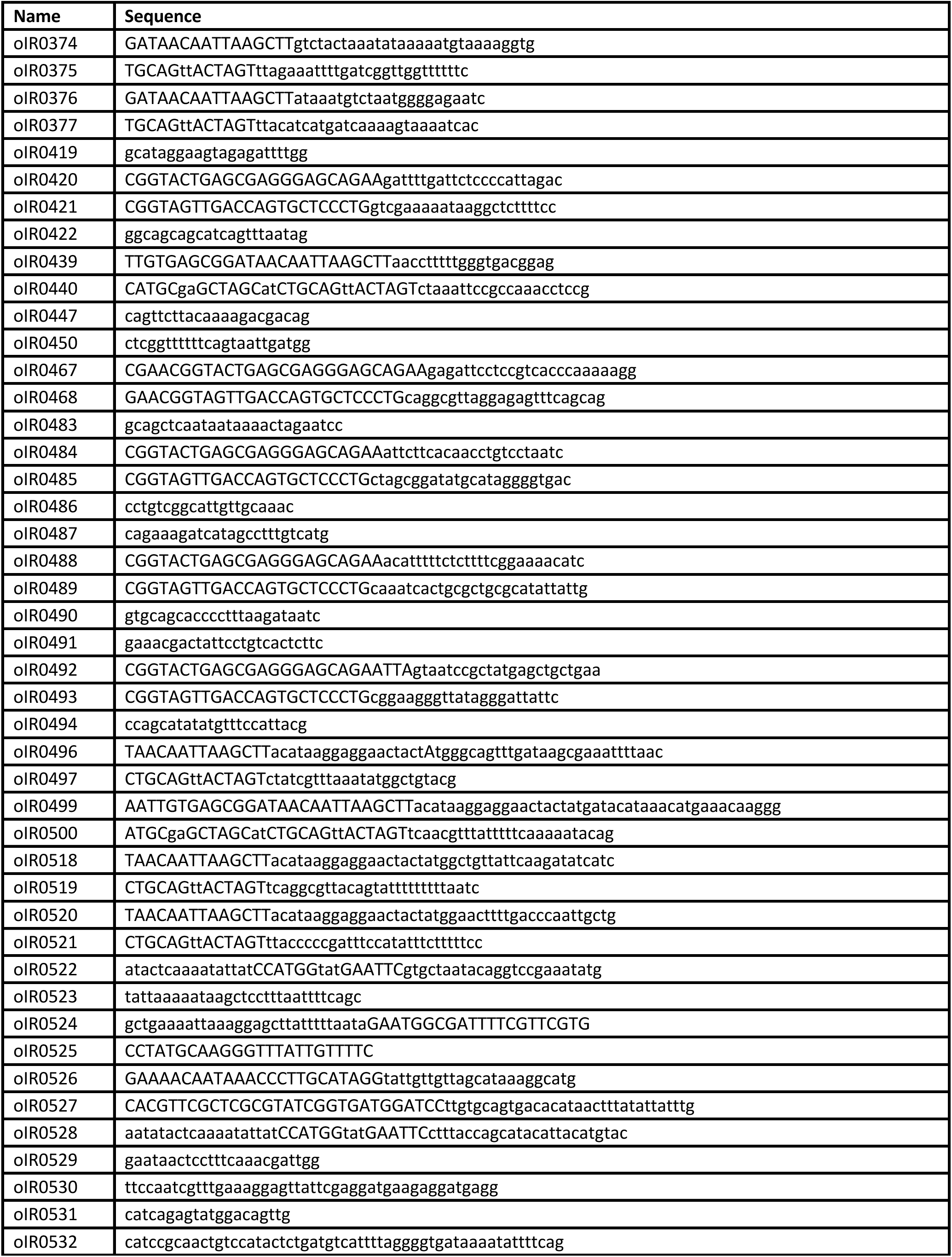
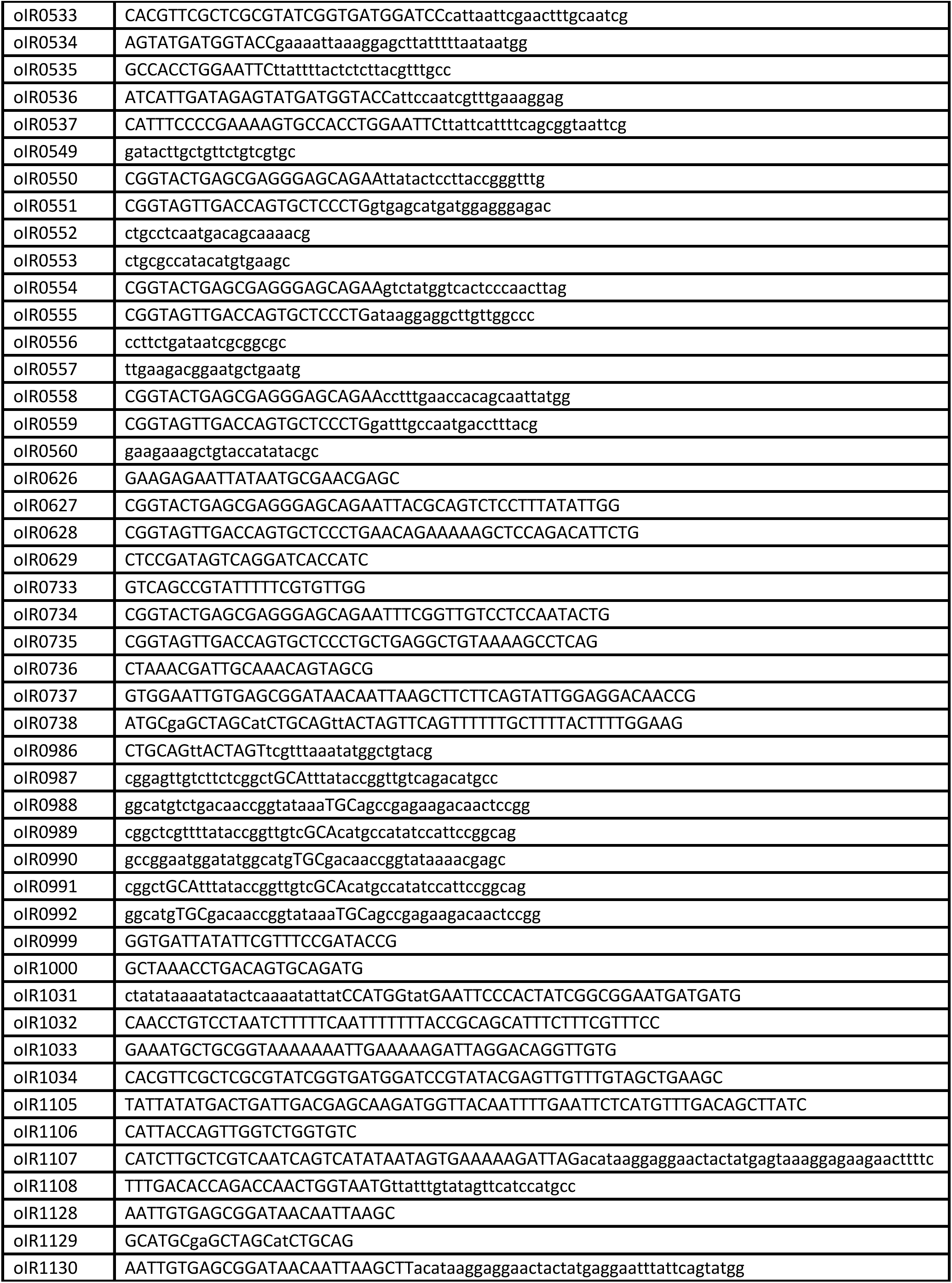
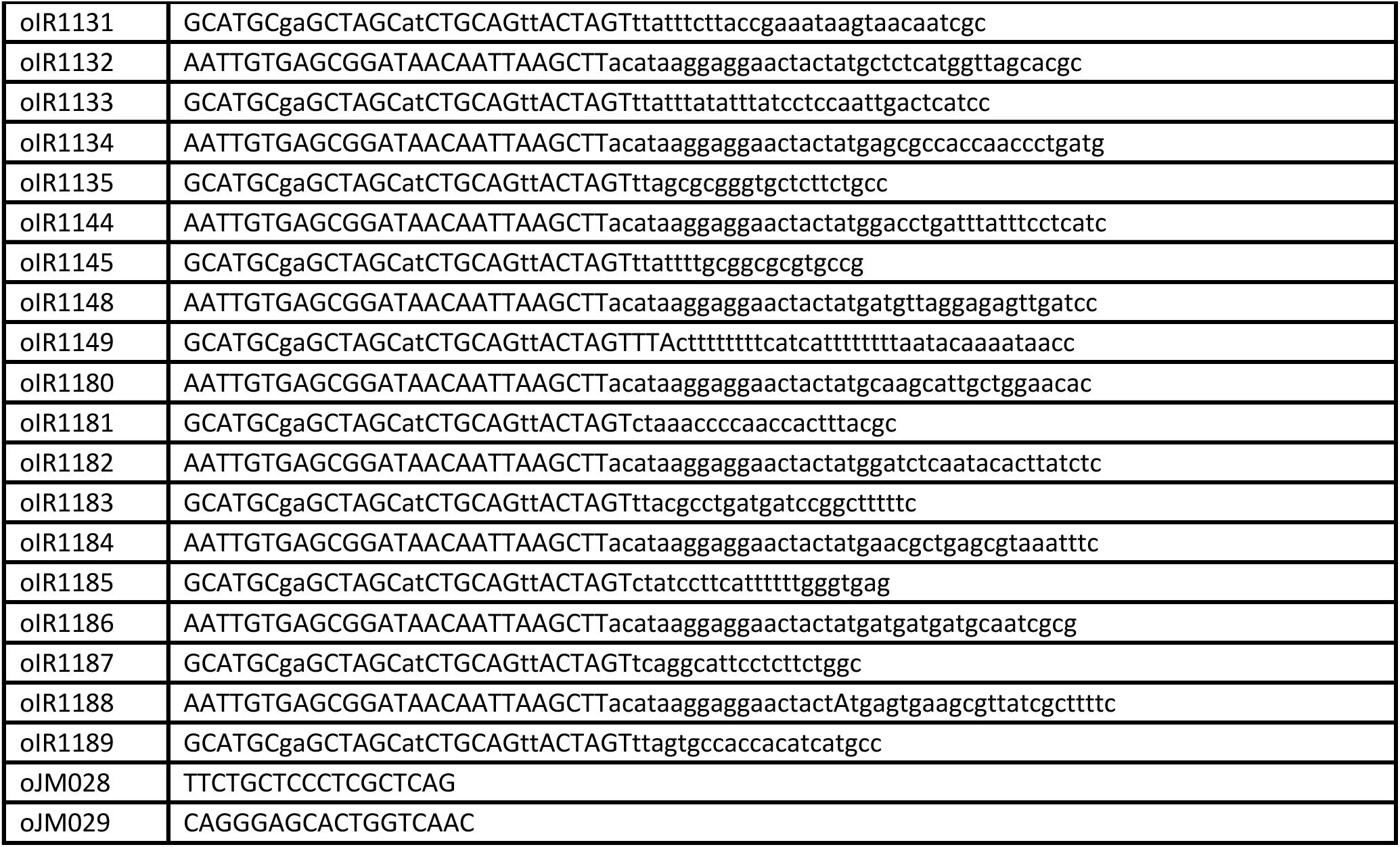
oligonucleotides used in this study.

**Table S6.**
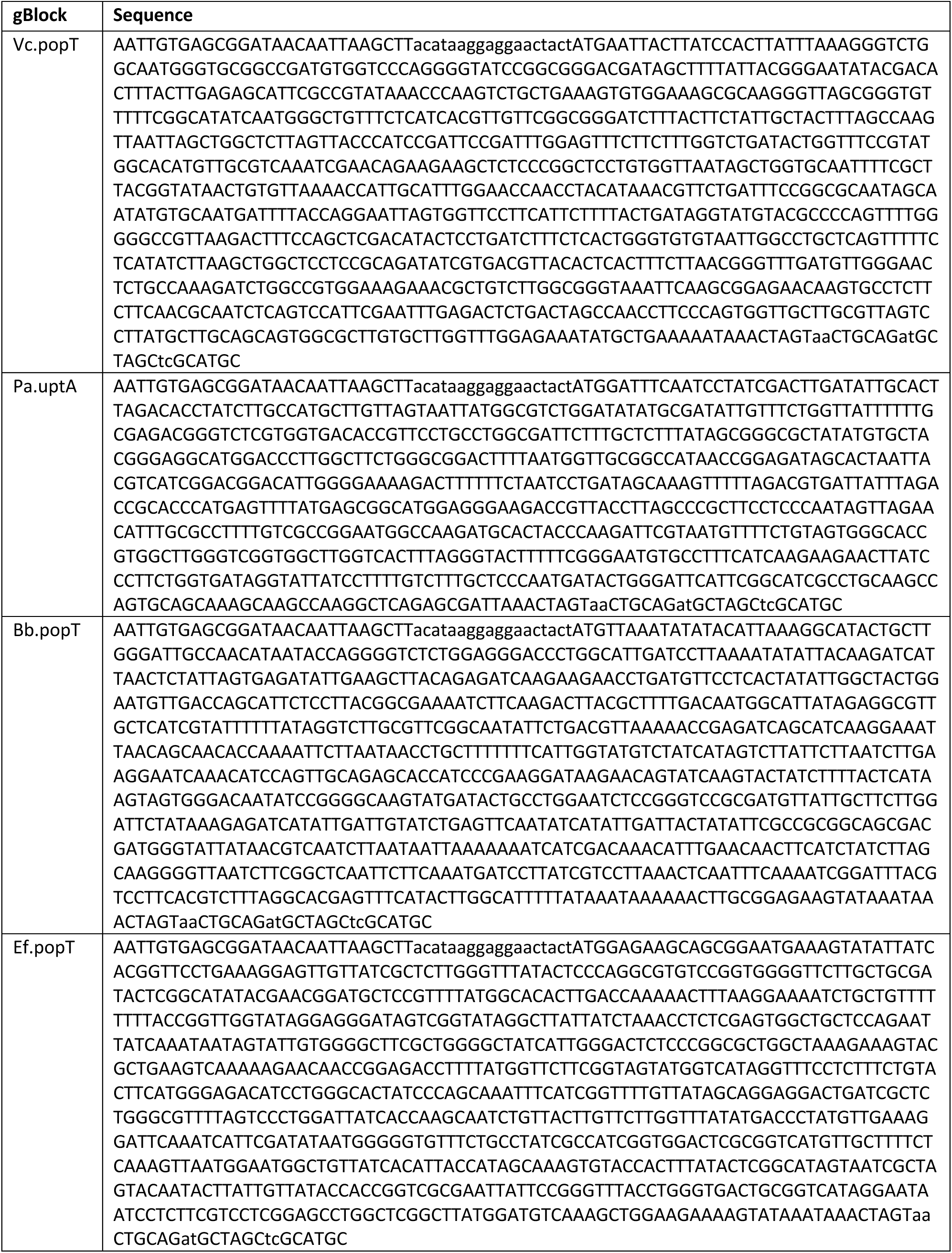
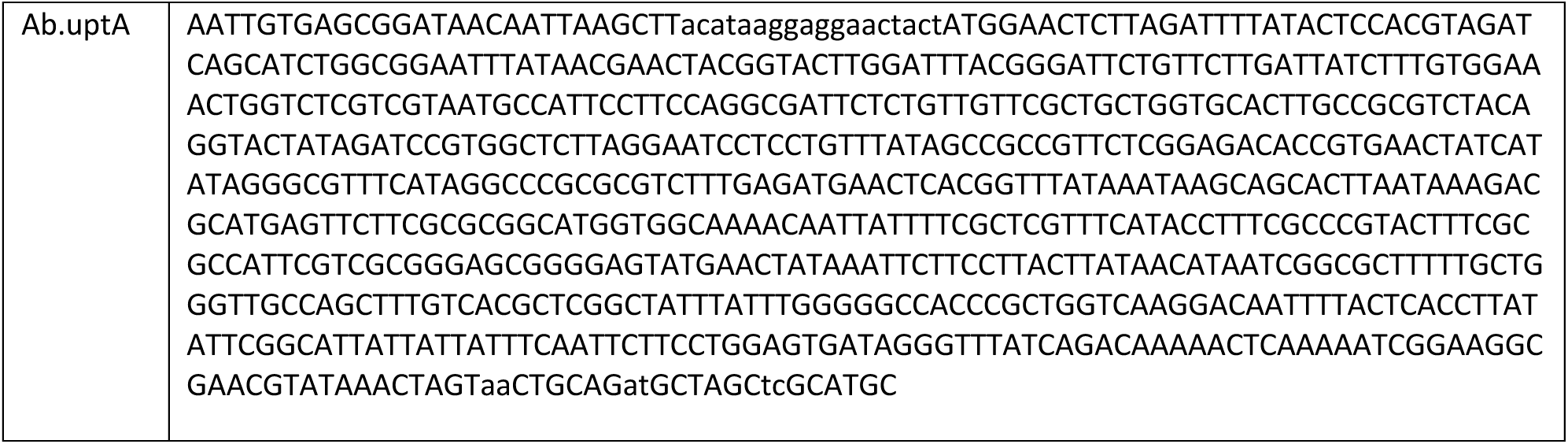
gBlocks used in this study.

## Supplemental Methods

### Materials and Methods

#### General methods and strain construction

All *Bacillus subtilis* strains were derived from the prototrophic strain PY79 (*1*). All *B. subtilis* experiments were performed at 37 °C with aeration in lysogeny broth (LB). Antibiotic concentrations used were: 100 µg/mL spectinomycin, 10 µg/mL kanamycin, 5 µg/mL chloramphenicol, 10 µg/mL tetracycline, 1 µg/mL erythromycin and 25 µg/mL lincomycin. Isopropyl ß-d-thiogalactopyranoside (IPTG) was used at a concentration of 500 µM unless indicated otherwise. All experiments with MX2401 or amphomycin used media supplemented with 25 µg/mL CaCl_2_. All *B. subtilis* strains were generated using the one-step competence method unless indicated otherwise. All *Staphylococcus aureus* strains were derived from RN4220. Cells were grown at 37 °C with aeration in Tryptic soy broth (TSB). Antibiotic concentrations used were: 7.5 µg/mL erythromycin, 50 µg/mL kanamycin, 450 µg/mL spectinomycin, 5 µg/mL chloramphenicol. Anhydrotetracycline (aTc) was used at a concentration of 100 ng/mL unless indicated otherwise. Transductions were performed using phage 80alpha. All strains, plasmids, oligonucleotides, and synthetic DNA used in this study can be found in Supplementary Tables S3, S4, S5, S6.

All MIC and spot-dilution assays presented were from one of three biological replicates with two or three technical replicates. All microscopy images were from one of at least two biological replicates.

#### Transposon mutagenesis and suppressor isolation

Transposon libraries were generated as described previously (*2*), libraries were generated using pIR242, an *E. coli* - *B. subtilis* shuttle vector containing a temperature-sensitive replicon for *B. subtilis*, the mariner-Himar1 transposase, and a spectinomycin resistance cassette followed by a strong outward-facing promoter (P*pen*) and flanked by inverted repeats recognized by the transposase. The pIR242 plasmid was separately transformed into and BIR769 or BIR672 and plated on LB agar supplemented with MLS and incubated at 30 °C. Transformants were inoculated into LB supplemented with spectinomycin and grown with aeration at 22 °C for 24 hours. Cultures were pooled and frozen with 15% glycerol. Aliquots of the frozen stocks were thawed and plated onto LB agar supplemented with spectinomycin, CaCl_2_ and 2.5 µg/mL MX2401 (BIR769) or 0.3 µg/mL MX2401 (BIR672). Plates were incubated at 42 °C overnight to select for transposon mutagenized cells that had lost the plasmid and provided resistance to MX2401.

#### Mapping transposon insertion sites

Sequencing and mapping of the transposon insertion sites was performed as described previously (*2, 3*). Briefly, transposon-mutagenized colonies that grew on LB agar supplemented with MX2401 were pooled, and genomic DNA extracted. DNA was digested with MmeI and ligated to adaptors. Transposon-chromosome junctions were amplified by PCR and the products were size selected using a 2% agarose gel. The Illumina MiSeq platform was used to sequence the library. Sequencing reads were mapped to the *B. subtilis* 168 chromosome (NCBI NC_00964.3).

#### Minimal inhibitory concentration (MIC) assays

Exponentially growing cultures of *B. subtilis* or *S. aureus* were back-diluted 1:1000 into 96-well microtiter plates containing the indicated concentrations of antibiotic and inducers. Plates were sealed with breathable membranes and grown with orbital shaking at 37 °C overnight. Plates were photographed after the overnight (∼16 hr) incubation.

#### Spot-dilution assays

Late-log cultures were normalized to OD600=1 and seven 10-fold serial dilutions were made. 5 µL of each dilution were spotted onto LB agar supplemented with or without IPTG or onto TSB agar supplemented with or without aTc. Plates were incubated at 37 °C overnight and photographed the next day.

#### Genomic Neighborhood Analysis

The Enzyme Similarity Tool (EFI-EST) from the Enzyme Function Initiative (*4*) (available at: https://efi.igb.illinois.edu/efi-est/) was used to generate sequence similarity networks (SSN) for the DedA and DUF368 protein families from the pfam entries SNARE_assoc (PF09335) and DUF368 (PF04018) respectively. Due to the large size of the SNARE_assoc family, the UniRef90 database was used. The SSNs were then used as inputs for performing genomic neighborhood analysis using the Enzyme Function Initiative Genome Neighborhood Tool (*4*) (EFI-GNT) available at: https://efi.igb.illinois.edu//efi-gnt/index.php. Alignment score cutoffs of 35% were used for both analyses. Gene neighborhood diagrams (GND) were generated to visualize the 10 nearest genes surrounding all *<u>dedA</u>* and *duf368* members.

An SSN was also generated for the subset of DUF368 members in Archaea, this dataset was used as an input for performing gene neighborhood analysis and the resulting gene neighborhood networks (GNN) were uploaded and visualized in cytoscape. Cytoscape was used to examine and quantify the genes most found within a 10 gene window of *duf368*.

#### MX2401-FL

MX2401 has a single primary amine (*5*) and drug optimization efforts have found that it can be modified without loss of potency. MX2401 was labeled at this position with CF488A using the Mix-n-Stain CF Dye Small Ligand Labeling Kit (Biotium #92350) following the manufacture’s instructions. Briefly, 10 µL (0.1 µmol) of MX2401 dissolved in DMSO was mixed with 2 µL of reaction buffer. The mixture was added to CF488A dye and briefly vortexed. The reaction was incubated for 30 minutes in the dark at room temperature. 2 µL of quenching buffer was added to the reaction, briefly vortexed and incubated for 5 minutes in the dark. DMSO was added to a final volume of 100 µL generating a 1 mM solution of MX2401-FL.

#### MX2401-FL labeling and fluorescence microscopy

Exponentially growing cultures of *S. aureus* were incubated for 20 minutes with the fluorescent D-amino acids (FDAAs) HADA (Tocris) or RADA (Tocris) at final concentrations of 100 µM.

Exponentially growing cultures of *Bacillus subtilis* and FDAA labelled *S. aureus* were collected by centrifugation at 7000 RPM for 2 min. Cells were washed once with 1xPBS (pH 7.4) and re-suspended in 1/25^th^ volume of 1xPBS. MX2401-FL was added to a final concentration of 25 µM and incubated for 30 seconds. Cells were washed with 1xPBS, resuspended in 1/25 volume of 1xPBS, and spotted onto 1.5% agarose pads containing growth medium.

Phase and fluorescence microscopy was performed with a Nikon Ti inverted microscope using a Plan Apo 100x/1.4 Oil Ph3 DM objective, a Lumencore SpectraX LED illumination system and an Andor Zyla 4.2 Plus sCMOS camera. Chroma ET filter cubes (#49000, 49002, 49003 and 49008) were used for imaging BFP/HADA, MX2401-FL, YFP and mCherry/RADA/Propidium Iodide respectively. Exposure time of 50 ms was used for HADA, RADA, Propidium Iodide and mCherry; 400 ms was used for BFP; 200 ms was used for MX2401-FL and 1 s was used for YFP. Images were acquired with Nikon elements 4.3 software and analyzed using ImageJ (version2.3).

#### Fluorescence microscopy quantification

ImageJ was used to quantify fluorescent intensities. A vegetatively expressed blue fluorescent protein (BFP) was used to identify cell boundaries. Intensity values from the YFP channel were extracted.

#### Structural model visualization

Alphafold2 predictions of PopT (SAOUHSC_00846) and UptA (YNGC_BACSU) were downloaded from the AlphaFold Protein Structure Database (available at: https://alphafold.ebi.ac.uk/). PyMOL was used to visualize the structural models and generate images. Reentrant helixes are highlighted in pink and blue. Arginine residues critical for *yngC* function are shown as red spheres.

#### Immunoblot analysis

Immunoblot analysis was performed as described previously (*6*). Briefly, 1mL of exponentially growing cells were normalized by OD600 and harvested by centrifugation (2 min at 7000 RPM). The cell pellet was resuspended in lysis buffer (20 mM Tris pH 7.0, 10 mM MgCl_2_, 1mM EDTA, 1 mg/mL lysozyme, 10 µg/mL DNase I, 100 µg/mL RNase A, 1 mM PMSF, 1 µg/mL leupeptin, 1 µg/mL pepstatin) and incubated at 37 °C for 15 minutes. An equal volume of sample buffer (0.25 M Tris pH 6.8, 4% SDS, 20% glycerol, 10 mM EDTA, 10% β-mercaptoethanol) was added to the lysis reactions and vortexed briefly to complete lysis. Proteins were separated by SDS-PAGE on 12.5% polyacrylamide gels, transferred onto Immobilon-P membranes (Millipore) by electrophoretic transfer and blocked with 5% Milk in phosphate buffered saline with 0.5% Tween-20 (PBS-T). The blocked membranes were probed with monoclonal anti-His (1: 4000) (Genscript) or polyclonal anti-SigA (1:10,000) (*7*) antibodies diluted into 3% BSA in PBS-T. Primary antibodies were detected using horseradish peroxidase-conjugated goat anti-rabbit IgG or goat anti-mouse IgG (BioRad) and the Super Signal chemiluminescence reagent as described by the manufacturer (Pierce).

#### Dendrograms

Dendrograms were made using AnnoTree (*8*). All entries of pf04018 (DUF368) were mapped to representative dendrograms of bacteria and archaea. Hits are shown as blue lines.

#### Multiple Sequence Alignment

Multiple sequence alignments were performed using Clustal Omega (*9*) and visualized with Espript(*10*).

#### Strain constructions

##### *B. subtilis* deletion mutants

Most *B. subtilis* deletion mutants were made by isothermal assembly (*11*) followed by direct transformation. The assembly reactions contained three PCR products: two PCR products containing ∼1500 base pairs upstream and downstream of the gene to be deleted, and a third PCR product containing an antibiotic resistance cassette. Antibiotic resistance cassettes with surrounding lox66/lox71 sites were amplified from pWX465(cat), pWX466(spec), pWX467(erm), pWX469(tet) and pWX470(kan) using the primers oJM028 and oJM029. The flanking regions for the respective deletions were amplified using PY79 genomic DNA as template and the following primer sets: *bceAB*(oIR626-629); *yngC* (oIR483-486); *ykoX*(oIR487-490); *ybfM*(oIR491-494); *yhjE*(oIR549-552); *yqeD* (oIR553-556); *ytxB*(oIR557-560); *uppS*(oIR447/oIR467,oIR468/oIR450); *ispH*(oIR733-736); *uppP*(oIR419-422).

The *bcrC* deletion was from the BKE collection and was backcrossed twice into PY79 and PCR confirmed.

##### Construction of the Δ6 mutant BIR712 [Δ*yngC*, Δ*ykoX*, Δ*ybfM*, Δ*yhjE*, Δ*yqeD*, Δ*ytxB*]

A Δ3 strain, BIR656 [*sacA::Pveg-mTagBFP (phleo), amyE::Pamj-yfp (cat), yngC::erm, ykoX::kan, ybfM::tet*] was made by successive transformations of isothermal assembly products to delete yngC, ykoX and ybfM with lox66/71 flanked erm, kan and tet resistance cassettes respectively. The antibiotic resistance cassettes were then looped out from the Δ3 strain using pDR244 (*3*). The remaining three deletions of yhjE::kan, yqeD::erm and ytxB::tet were made by successive rounds of isothermal assembly and transformation to create BIR712 (*sacA::Pveg- mTagBFP (phleo), amyE::Pamj-YFP (cat), yngC::lox72, ykoX::lox72, ybfM::lox72, yhjE::kan, yqeD::erm, ytxB::tet)*.

##### Construction of the P*yngC*(ΔP*_sigM_*) mutant by allelic exchange

pIR361 [Pmad3.1-PyngC(ΔPsigM)(erm)(amp)] was passaged through a recA+ *E. coli* strain (AB1157) and transformed into PY79. A transformant obtained at 37°C was grown overnight at 22°C and then serial dilutions plated on LB agar at 37°C. Single colonies were streaked onto LB and LB+MLS to identify strains that had looped out the integrated plasmid. The yngC promoter was PCR amplified from MLS(S) strains and sequenced confirm the promoter deletion.

##### Construction of *yngC* point mutations

Point mutations in yngC-his10 were made by isothermal assembly and direct transformation into *B. subtilis*. Two DNA fragments were amplified using the genomic DNA of BIR1271 [ycgO-Phyperspank-yngC-his10-spec-ycgO] as a template using oligos flanking the upstream and downstream homology arms (oIR999 and oIR1000) and mutation specific primers (R112A = oIR987/oIR988, R118A = oIR989/oIR990, R112A,R118A = oIR991/oIR992). The two resulting amplification products were purified and added to the isothermal assembly reaction followed by direct transformation into BIR648. All mutants were confirmed by sequencing.

##### Construction of deletion mutants of *uptA*(*Sa*) and *popT*(*Sa*) in *S. aureus*

BIR683 [Δ02816::spec], BIR688 [Δ00846::kan] were made by allelic replacement of coding regions with antibiotic resistance cassettes using a loop-in-loop-out approach as has been previously described (*12*) with minor modifications. Briefly, pMad based plasmids (pIR237[pMAD3.1-02816::spec(erm)(amp)] and pIR238 [pMAD3.1-00846::kan(erm)(amp)] were electroporated into ATP001 (RN4220 WT) and selected on TSB agar + erm at 30C. Single colonies were inoculated in TSB + erm for 24 hours at 30C with two 1/100 back-dilutions. Serial dilutions were made of the cultures and aliquots were plated on TSA + erm at 42°C to lose unintegrated plasmids. Colonies were inspected for the expression of mScarlet which is constitutively expressed from the plasmid. Pink colonies, indicating plasmid integration, were selected and inoculated in TSB and grown without antibiotics at 30C for 24 hours with two 1/100 back-dilutions. Serial dilutions were made of the cultures and aliquots were plated on TSA at 30C. White colonies were streaked on TSA, TSA + erm and (TSA + spec or kan) to confirm the integration of the deletion cassette and loss of plasmid. The deletion mutants were confirmed by PCR with primers flanking the target gene.

##### Construction of complementation strains

BIR691 [attB::Ptet-02816 (cat)] and BIR695 [attB::Ptet-00846 (cat)] were made by electroporating pIR239 [attB::Ptet-02816 (cat)(amp)] or pIR240 [attB::Ptet-00846 (cat)(amp)] into a L54a integrase expressing strain RN4220 (pTP044) and selecting on TSA + Cm at 42°C to lose the integrase expressing plasmid. PIR239 and pIR240 were integrated into the attB (L54a) site within the *geh* gene. The correct integration was confirmed by PCR.

BIR715 [Δ02816::spec, Δ00846::kan] was constructed by phage transduction of the 846::kan deletion into BIR683[2816::spec] using phage 80alpha. PCR was performed to confirm the transduction with PCR primers flanking the deleted genes.

BIR1279 [[Δ02816::spec, Δ00846::kan, attB::Ptet-02816 (cat)], BIR1280 [[Δ02816::spec, Δ00846::kan, attB::Ptet-00846 (cat)] were constructed by successive rounds of phage transduction of the 2816::spec and 846::kan markers respectively. The gene deletions were confirmed by PCR.

#### Plasmid Constructions

**pIR233 [ycgO::Phyperspank-bcrC (spec) (amp)]**

pIR233 was generated in a two-piece ligation with PCR product containing the bcrC gene (amplified from PY79 gDNA with oIR374 and oIR375) and pCB090 [ycgO::Phyperspank(spec)] digested with HindIII and SpeI.

**pIR234 [ycgO::Phyperspank-uppP (spec)(amp)]**

pIR234 was generated in a two-piece ligation with PCR product containing the uppP gene (amplified from PY79 gDNA with oIR376 and oIR377) and pCB090 [ycgO::Phyperspank(spec)] digested with HindIII and SpeI.

**pIR224 [ycgO::Phyperspank-yngC (spec) (amp)]**

pIR224 was generated in a two-piece ligation with PCR product containing the yngC gene (amplified from PY79 gDNA with oIR496 and oIR497) and pCB090 [ycgO::Phyperspank(spec)] digested with HindIII and SpeI.

**pIR226 [ycgO:: Phyperspank-ykoX (spec) (amp)]**

pIR226 was generated in a two-piece isothermal assembly reaction with PCR product containing the ykoX gene (amplified from PY79 gDNA with oIR499 and oIR500) and pCB090 [ycgO::Phyperspank(spec)] digested with HIndIII and SpeI.

**pIR239 [attB::Ptet-02816 (cat) (amp)]**

pIR239 was generated in a two-piece ligation with PCR product containing the SAOUHSC_02816 gene (amplified from RN4220 gDNA with oIR534 and oIR535) and pTB005[attB::Ptet-02816] digested with KpnI and EcoRI.

**pIR240 [attB::Ptet-00846 (cat) (amp)]**

pIR240 was generated in a two-piece isothermal assembly reaction with PCR product containing the SAOUHSC_00846 gene (amplified from RN4220 gDNA with oIR536 and oIR537) and pTB005[attB::Ptet-02816] digested with KpnI and EcoRI.

**pIR237 [Pmad3.1-2816::spec (amp)]**

pIR237 was generated in a four-piece isothermal assembly reaction with PCR products flanking the SAOUHSC_02816 gene (amplified from RN4220 gDNA with oIR522/oIR523 and oIR526/oIR527), a PCR product encoding the spectinomycin resistance cassette (amplified from pWX466 with oIR524/oIR525) and pMR091[Pmad3.1 (amp)] digested with BamHI and EcoRI.

**pIR238 [Pmad3.1-846::kan (amp)]**

pIR237 was generated in a four-piece isothermal assembly reaction with PCR products flanking the SAOUHSC_00846 gene (amplified from RN4220 gDNA with oIR528/oIR529 and oIR532/oIR533), a PCR product encoding the kanamycin resistance cassette (amplified from pWX470 with oIR530/oIR531) and pMR091[Pmad3.1 (amp)] digested with BamHI and EcoRI.

**pIR400 [ycgO::Phyperspank-popT(Sp) (spec)(amp)]**

pIR226 was generated in a two-piece isothermal assembly reaction with PCR product containing the SPD_0872 gene (amplified from S.pneumoniae D39 gDNA with oIR1132 and oIR1133) and pCB090 [ycgO::Phyperspank(spec)] digested with HindIII and SpeI.

**pIR388 [ycgO::Phyperspank-popT(Vc) (spec)(amp)]**

pIR226 was generated in a two-piece isothermal assembly reaction with PCR product containing the MS6_A0029 gene of V. cholerae C6706 (amplified from gBlock Vc.popT with oIR1128 and oIR1129) and pCB090 [ycgO::Phyperspank(spec)] digested with HindIII and SpeI.

**pIR399 [ycgO::Phyperspank-popT(Bc) (spec)(amp)]**

pIR226 was generated in a two-piece isothermal assembly reaction with PCR product containing the BC5158 gene of B. cereus atcc14579 (amplified from gDNA with oIR1130 and oIR1131) and pCB090 [ycgO::Phyperspank(spec)] digested with HindIII and SpeI.

**pIR397 [ycgO::Phyperspank-uptA(Pa) (spec)(amp)]**

pIR397 was generated in a two-piece isothermal assembly reaction with PCR product containing the PA4029 gene of P.aeruginosa PAO1 (amplified from gBlock Pa.uptA with oIR1128 and oIR1129) and pCB090 [ycgO::Phyperspank(spec)] digested with HindIII and SpeI.

**pIR393 [ycgO::Phyperspank-popT(Bb) (spec)(amp)]**

pIR393 was generated in a two-piece isothermal assembly reaction with PCR product containing the BBUBOL26_RS01045 gene of B. burgdorferi (amplified from gBlock Bb.popT with oIR1128 and oIR1129) and pCB090 [ycgO::Phyperspank(spec)] digested with HindIII and SpeI.

**pIR405 [ycgO::Phyperspank-uptA(Bc) (spec)(amp)]**

pIR405 was generated in a two-piece isothermal assembly reaction with PCR product containing the BC5040 gene of B. cereus atcc14579 (amplified from gDNA with oIR1148 and oIR1149) and pCB090 [ycgO::Phyperspank(spec)] digested with HindIII and SpeI.

**pIR411 [ycgO::Phyperspank-popT(Ef) (spec)(amp)]**

pIR411 was generated in a two-piece isothermal assembly reaction with PCR product containing the gene of E. faecium 13.SD.W.09 (amplified from gBlock Ef.popT with oIR1128 and oIR1129) and pCB090 [ycgO::Phyperspank(spec)] digested with HindIII and SpeI.

**pIR401 [ycgO::Phyperspank-popT(Cg) (spec)(amp)]**

pIR401 was generated in a two-piece isothermal assembly reaction with PCR product containing the CGP_RS06655 gene of C. glutamicum MB001 (amplified from gDNA with oIR1134 and oIR1135) and pCB090 [ycgO::Phyperspank(spec)] digested with HindIII and SpeI.

**pIR211 [ycgO::Pspank-uppS (spec)(amp)]**

pIR211 was generated in a two-piece isothermal assembly reaction with PCR product containing the uppS gene (amplified from PY79 gDNA with oIR439 and oIR440) and pCB084 [ycgO::Pspank(spec)] digested with HindIII and SpeI.

**pIR278 [ycgO::Pspank-ispH (spec)(amp)]**

pIR278 was generated in a two-piece isothermal assembly reaction with PCR product containing the ispH gene (amplified from PY79 gDNA with oIR737 and oIR738) and pCB084 [ycgO::Pspank(spec)] digested with HindIII and SpeI.

**pIR235 [ycgO::Phyperspank-yghB (spec)(amp)]**

pIR235 was generated in a two-piece ligation with PCR product containing the yghB gene (amplified from MG1655 gDNA with oIR518 and oIR519) and pCB090 [ycgO::Phyperspank(spec)] digested with HindIII and SpeI. The resulting plasmid was sequence confirmed.

**pIR236 [ycgO::Phyperspank-yqjA (spec)(amp)]**

pIR236 was generated in a two-piece ligation with PCR product containing the yqjA gene (amplified from MG1655 gDNA with oIR520 and oIR521) and pCB090 [ycgO::Phyperspank(spec)] digested with HindIII and SpeI.

**pIR403[ycgO::Phyperspank-dedA (spec)(amp)]**

pIR403 was generated in a two-piece isothermal assembly reaction with PCR product containing the dedA gene (amplified from MG1655 gDNA with oIR1144 and oIR1145) and pCB090 [ycgO::Phyperspank(spec)] digested with HindIII and SpeI.

**pIR406 [ycgO::Phyperspank-yabI (spec)(amp)]**

pIR406 was generated in a two-piece isothermal assembly reaction with PCR product containing the yabI gene (amplified from MG1655 gDNA with oIR1180 and oIR1181) and pCB090 [ycgO::Phyperspank(spec)] digested with HindIII and SpeI.

**pIR407 [ycgO::Phyperspank-yohD (spec)(amp)]**

pIR407 was generated in a two-piece isothermal assembly reaction with PCR product containing the yohD gene (amplified from MG1655 gDNA with oIR1182 and oIR1183) and pCB090 [ycgO::Phyperspank(spec)] digested with HindIII and SpeI.

**pIR408 [ycgO::Phyperspank-ydjX (spec)(amp)]**

pIR408 was generated in a two-piece isothermal assembly reaction with PCR product containing the ydjX gene (amplified from MG1655 gDNA with oIR1184 and oIR1185) and pCB090 [ycgO::Phyperspank(spec)] digested with HindIII and SpeI.

**pIR409 [ycgO::Phyperspank-ydjZ (spec)(amp)]**

pIR408 was generated in a two-piece isothermal assembly reaction with PCR product containing the ydjZ gene (amplified from MG1655 gDNA with oIR1186 and oIR1187) and pCB090 [ycgO::Phyperspank(spec)] digested with HindIII and SpeI.

**pIR410 [ycgO::Phyperspank-yqaA (spec)(amp)]**

pIR408 was generated in a two-piece isothermal assembly reaction with PCR product containing the yqaA gene (amplified from MG1655 gDNA with oIR1188 and oIR1189) and pCB090 [ycgO::Phyperspank(spec)] digested with HindIII and SpeI.

**pIR385 [amyE::PyngC-yfp (cat)(amp)]**

pIR385 was generated in a three-piece isothermal assembly reaction with PCR product containing the yfp gene with the yngC promoter in the primer overhang (amplified from pAM155 with oIR1105 and oIR1106 ) and pDG364 [amyE::cat] (PCR linearized with oIR1107 and oIR1108).

**pIR361 [Pmad3.1-PyngC(ΔPsigM) (erm)(amp)]**

pIR361 was generated in a three-piece isothermal assembly reaction with PCR products flanking the sigM promoter of yngC (oIR1031/oIR1032 and oIR1033/oIR1034) and pMR091 digested with BamHI and EcoRI.

**pIR392 [ycgO::Phyperspank-uptA(Ab) (spec)(amp)]**

pIR392 was generated in a two-piece isothermal assembly reaction with PCR product containing the AB17975 gene of A. baumanni (amplified from gBlock Ab.uptA with oIR1128 and oIR1129) and pCB090 [ycgO::Phyperspank(spec)] digested with HindIII and SpeI.

**pIR352 [ycgO::Phyperspank-yngC-his10 (spec)(amp)]**

pIR352 was generated in a two-piece ligation with PCR product containing the yngC gene (amplified from PY79 gDNA with oIR496 and oIR986) and pIR301 [ycgO::Phyperspank-MCS-his10(spec)] digested with HIndIII and SpeI.

All plasmids were sequence confirmed. All gene fusions to the Phyperspank promoter contained a synthetic optimized ribosome binding. All gene fusions to the Pspank promoter contained the native ribosome binding site.

## Notes

### Competing Interest Statement

The authors have declared no competing interest.

